# Entanglement of AGE-RAGE axis in cardiac pathosis

**DOI:** 10.1101/2023.07.23.550244

**Authors:** Rufaida Wasim, Tarique Mahmood, Mohammed Haris Siddiqui, Tanveer A. Wani, Seema Zargar, Aditya Singh, Saad Mohammed, Farogh Ahsan, Muhammad Wahajuddin

## Abstract

Cardiovascular diseases are the major cause of death globally. Acute coronary syndrome is one of numerous cardiovascular illnesses, including advanced glycation end products (AGEs), which play an instrumental part in their development and progression. A substance with multiple pleiotropic characteristics is rosuvastatin. This study examined the cardioprotective effects of rosuvastatin in isoproterenol-induced myocardial injury, as well as the alterations in advanced glycation end products and their roles in cardiac damage. Rosuvastatin (10 mg/kg, orally) was given to male rats daily for 4 weeks, and on the 29th and 30th days, isoproterenol (85 mg/kg, subcutaneously) was administered to cause cardiac damage. Rats were euthanised on the 31st day, and various samples were collected for examination. Administration of isoproterenol increased cardiac mass, levels of cardiac damage markers, lipid oxidation, and collagen content in the heart. Additionally, it reduced the activities of SOD, CAT, GST, GR, and all other antioxidants. Additionally, isoproterenol raises levels of inflammatory markers including TNF-α and IL-6. It has been observed that advanced glycation end products rise along with heart injury. The AGE-RAGE cascade also messes with the injured heart’s echocardiogram. Additionally, histopathological alterations were noticed. According to the study, rosuvastatin has cardioprotective effects on the experimental model, which were supported by an array of physical, biochemical, and histological characteristics.

**Summary:** This study examined the cardioprotective effects of rosuvastatin in isoproterenol-induced myocardial injury, as well as the alterations in advanced glycation end products and their roles in cardiac damage. According to the study, rosuvastatin has cardioprotective effects on the experimental model, which were supported by a number of physiological, biochemical, and histological characteristics. By showing how the functional AGE/RAGE axis is inhibited following rosuvastatin medication, this work provides an etiologic theory involving rosuvastatin therapy in heart injury. These findings may also have practical significance since they highlight the intriguing possibility that rosuvastatin-induced modulation of AGE-RAGE signalling may represent a unique therapeutic approach for the prevention and treatment of other cardiovascular diseases.

## 1. Introduction

Acute myocardial infarction (AMI) generally develops by atherosclerotic-mediated blockage of the coronary arteries, which inhibits nutrient convey to cardiomyocytes, expedites cell death and impede cardiac function. The myocardial ischemia stimulate inflammatory signalling, escorted by the action of immune cells, yielding reactive oxygen species, and molecular damage [1].

Advanced glycation end products (AGEs) constitute a heterogeneous aggregation of compounds inferred from the interplay of reducing sugars and/or other α-carbonyl compounds with amino groups [2] within proteins, lipids, and nucleic acids [3]. The impetus of AGE formation is Millard reaction. The first reversible step encompass condensation of reducing sugars with a free amino group [4]. The acquired Schiff’s base (aldimin) endures Amadori rearrangement to form ketosamine structure. Then the Amadori product undergoes further rearrangements, oxidations and elimination, and finally forms AGE [2]. This reaction is fast and depends on the concentration of available carbohydrates, being enhanced in hyperglycemia [3, 5, 6].

The deposition of glycated products is due to both an increased AGE production and an impaired degradation of modified proteins. Glycated intracellular proteins are cleaved slowly by the ubiquitin-proteasome system [4]. Partial proteolysis and/or intracellular capture mediated through AGER-1 receptor [2] are among the involved mechanisms. AGE degradation in macrophages generates soluble low molecular peptides, named second generation AGEs [2].

Ott et al. suggested that AGE accumulation is possible in long-lived proteins, e.g., collagen of extracellular matrix [4]. Some factors, such as the concentration and reactivity of glucose, the availability of precursors of AGE [4], and oxidative stress [6] can accelerate AGE formation, resulting in structural and/or functional changes affecting short-lived substrates [2].

AGEs significantly contribute to tissue and organ dysfunction [4], triggering oxidative stress, inflammation and apoptosis due to stimulation of specific receptors (RAGE) [2]. The mechanisms underlying the biological effects are accounted for the following hypotheses: *glycation modifies the structure and inhibits the function of proteins [5]; **glycation induces protein cross-linking and leads to tissue stiffness [5,7]; ***soluble AGEs bind to RAGE receptors and activate the intracellular signalling pathways (p21ras/MAPK, NADPH-oxidase/ ROS/ NF-kB; Jak/Stat), resulting in the expression of inflammatory cytokines (IL6, TNFα) and tissue inflammation [4].

AGEs also alter lipoproteins contributing to the development of atherosclerosis [2, 6, 7]. Both glycated and oxidized LDL and HDL reduce glutathione peroxidase-1 activity, and induce production of reactive oxygen species (ROS), endoplasmic reticulum stress, and apoptosis [2]. In 2014 Stirban et al. established that glycation and oxidation of LDL amplify their atherogenic effects, whereas oxidized and glycated HDL partially loses its antioxidant properties and protective role [2].

Experimental research has shown that AGEs cause intricate vascular lesions. Circulating AGEs enhance vascular permeability, cause atherosclerotic lesions of the aorta, and cause the cross-linking of connective tissue proteins in the arterial wall [2].

Clinical studies with a small sample size have demonstrated AGE involvement in acute coronary syndrome (ACS). In patients with ischemic heart disorders, AGE values correlate with the amount of coronary lesions, according to a study by Kiuchi et al [8]. In non-diabetic patients with ACS, Raposeiras Roubin et al. hypothesised the value of plasma AGE levels as long-term predictors of mortality, reinfarction, and risk of developing heart failure [7].

Statins, which are 3-hydroxy-3-methylglutaryl-coenzyme A reductase inhibitors, are often used to treat coronary heart disease due to their capacity to lower cholesterol [9]. Statins also stabilise plaque, lower inflammation, and guard against endothelial dysfunction, among other good pleiotropic effects [10]. Through the HMGB1/TLR4 pathway, statins also protect endothelial cells from injury caused by ischemia-reperfusion [11,12). But additional information is required to understand how statins might impact the HMGB1/RAGE/NF-ҝB pathway and the production of inflammatory chemicals. Rosuvastatin is a statin of the most recent generation and an inhibitor of HMG CoA reductase. It displays pleotropic effects without inhibiting HMG CoA reductase. A few of these include enhanced endothelial function, anti-inflammatory, anti-thrombotic, and antioxidant capabilities, as well as a decreased risk of cardiovascular death and events. It significantly affects a variety of cellular processes and mitigates AGE-RAGE’s negative consequences. Therefore, by improving the AGE-RAGE axis, it can close the research gap. Major cardiac conditions including myocardial ischaemia and associated illnesses may be helped by rosuvastatin. This study demonstrates the clinical relationship between the AGE-RAGE axis and heart injury. This study also highlights rosuvastatin’s preventive effects against heart diseases. This study examined the cardioprotective effects of rosuvastatin in isoproterenol-induced myocardial injury, as well as the alterations in advanced glycation end products and their roles in cardiac damage.

## 2. Materials and methods

### 2.1. Animal

The study included 24 adult male Sprague Dawley rats weighing 120-180 g. The animals were kept in Animal house, Faculty of Pharmacy Integral University Lucknow under standard conditions: in polypropylene cages, housed at 24 ± 2°C, alternating light/ dark cycle every 12 hours, daily-performed examination and cleaning of animal cages, and free access to food and water. The animals were fasted for 12 hours prior to the sacrifice. The animals were maintained and used in accordance with the Institutional Animal Ethics Committee.

### 2.2. Ethical Approval

The entire experiment was conducted according to the guidelines of CPCSEA. The research protocol was affirmed by Institutional Animal Ethical Committee (IAEC) with approval number (IU/IAEC/21/09), (Reg no.1213/PO/Re/S/08/CPCSEA, 5 June 2008) of faculty of pharmacy, Integral University, Lucknow (U.P.) India.

### 2.3. Treatment Protocol

Rats were weighed (150–180 g) and randomly divided into four groups (n = 6) with six rats in each group as described below. Prior to dosing, they were housed in their separate cages for seven days to allow them to acclimatise to the circumstances of the study facility and receive individual identification markings.

Group I Normal Control group (NCG): Rats were administered with normal saline 10mg/kg p.o for 28 days.

Group II Isoproterenol group (ISG): Isoproterenol (85 mg/kg/day, s.c.) twice at an interim of 24 h on 29th and 30th day.

Group III Treatment group (TG): Rosuvastatin (10 mg/kg b.w.) dissolved in normal saline and administered per orally (p.o.) by intubation method once a day for 28 days and isoproterenol (85 mg/kg/day, s.c.) twice at an interim of 24 h on 29th and 30th day.

Group IV *Per se* group (PSG): Rosuvastatin (10 mg/kg b.w.) dissolved in normal saline and administered per orally (p.o.) by intubation method once a day for 28 days.

### 2.4. Serum Preparation

Blood was collected and allowed to clot for around 40 minutes at room temperature in a dry test tube. Centrifugation at about 5000 rpm for 10 min was used to separate the serum. After centrifugation, serum was carefully extracted using a micropipette without destroying any of the tube’s cellular components. The cardiac markers assay was conducted using serum samples.

### 2.5. Tissue homogenate preparation

The 0.3 g of cardiac tissue was homogenised in 3 mL of 0.25 M sucrose buffer (pH 7.4) on ice. The resulting homogenate was centrifuged after being treated with 30 L of Triton X-100 and chilled for 30 minutes. glucose and AGE levels in a myocardial homogenate that was spun for ten minutes at 3000 rpm and 4°C. Until analysis, the supernatant was kept at −40°C. The collected blood samples were centrifuged for 10 minutes at 1500 rpm after being allowed to coagulate in test tubes for 30 minutes. Until analysis, serum was kept in Eppendorfs at −40°C.

### 2.6. Plasma Preparation

The plasma was separated from the blood after it had been collected in a centrifuge tube that had been heparinized and centrifuged for 10 minutes at 4000 rpm. During handling, the temperature of the plasma was kept between 2 and 8 C. The residual plasma was divided into 0.5 mL aliquots and kept at 20 °C; the fresh separated plasma was used for the examination of several biochemical parameters [13].

### 2.7. Measurement of Heart:Body Weight Ratio

After the animals were euthanised, their body weight was measured and recorded. Rats were pinned to the board with their extended extremities (inside hands and foot side) while lying on their backs. The mouse was washed or moistened with ethanol at a 70 percent concentration to remove hair and dander. After performing aortic root dissection directly above the aortic valves and superior vena cava dissection above the atria, the heart was removed. Forceps were used to carefully remove the excised heart’s mediastinal fat pads. By pressing the heart on a Kim wipe (via an absorbent pad) or surgical compress, blood from the heart was removed. The heart was dried entirely by repeating this cycle. The dried heart was weighed and documented after which the heart was fixed.[14].

### 2.8. Echocardiography measurements

The rats’ cardiac function was evaluated using echocardiography. Through intraperitoneal injection, the rats were anesthetized using chloral hydrate (5%, 0.7 ml/100 g, or 350 mg/kg). Using a Vivid 7 echocardiography device (GE Healthcare Life Sciences, Chalfont, UK) and a 12-MHz linear transducer, pictures were taken. The level of the papillary muscle was used to get a two-dimensional short-axis image of the left ventricle and to capture two-dimensional targeted M-mode tracings. Left ventricular ejection fraction (LVEF), left ventricular end-diastolic pressure (LVEDP), left ventricular fractional shortening (LVFS), systolic left ventricular internal dimension (LVIDs), and diastolic left ventricular internal dimension (LVIDd), were the detecting indicators.

### 2.9. Oxidative Stress Parameter

The Thiobarbituric Acid (TBA) Test Technique of Buege and Aust, (1978) was used to measure the malondialdehyde [15]. TCA-TBA-HCl reagent and 0.4 mL of serum were combined in a brief amount of time. mixed and left in the water bath for 10 minutes. A freshly prepared 1 N NaOH solution was added to 1 mL after cooling. At 535 nm, the pink color’s visibility was assessed in comparison to a white background. It was determined how much lipid peroxide was present using the method reported by Ohkawa et al. 1979 [16]. I1 mL of 10% cardiac tissue homogenate underwent 10,000 rpm for 10 min of centrifugation. This was then centrifuged for 15 minutes at 3000 rpm for 15 minutes while being maintained at 80 C for a volume of 0.5 mL of 0.8% TBA and 0.5 mL of 30% TCA. The absorbance was calculated at 540 nm. The lipid hydroperoxide in the tissues and plasma was measured using Jiang et al.’s estimation method in 1992. In brief, 0.2 mL of sample was combined with 1.8 mL of Fox reagent, which was then incubated for 30 minutes at 37 °C before being centrifuged for 10 minutes at 10,000 rpm. Absorbance was found at 540 nm [17]. Conjugated dienes were studied using the technique outlined by Rao and Recknagel in 1968. In a nutshell, 0.1 mL of tissue homogenate or plasma was mixed with 5.0 mL of chloroform-methanol (2:1 v/v) reagent. Next, the mixture was centrifuged for five minutes. Centrifugation was used to remove the bottom layer and evaporate it to dryness. The absorbance was measured at 233 nm following the addition of 1.5 mL of cyclohexane [18].

### 2.10. Measurement of Antioxidant Parameters

The level of ascorbic acid in the serum or plasma was assessed as a non-enzymatic antioxidants measure. In a nutshell, the supernatant’s absorbance at 700 nm was evaluated in comparison to a blank solution made from 2 mL of distilled water and 2 mL of a colour reagent. To assess serum vitamin E levels, as demonstrated by Baker and Frank (1968). In [19], the colorimetric approach was utilised. A test tube containing 1.5 mL of serum, 1.5 mL of ethanol, and 1.5 mL of xylene was thoroughly mixed before being centrifuged at 5000 rpm for 10 min. 1 mL of the xylene layer was separated before using the dipyridyl reagent. To assess serum vitamin E levels, as demonstrated by Baker and Frank (1968). In [19], the colorimetric approach was utilised. A test tube containing 1.5 mL of serum, 1.5 mL of ethanol, and 1.5 mL of xylene was thoroughly mixed before being centrifuged at 5000 rpm for 10 min. 1 mL of the xylene layer was separated before using the dipyridyl reagent.

Non-enzymatic antioxidants marker (heart): Ascorbic acid was measured using the titration method explained by Sadasivam and Manickam (1992) [21]. In brief, oxalic acid was used to homogenise 50 mg of fresh tissue with 5 mL before filtering and titrating with dichlorophenol indophenols. It was calculated how much dye was consumed. The procedure Varley H (1976) [22] outlined is used to quantify-tocopherol (Vitamin E). A 1.5 mL mixture of cardiac tissue extract and 1.5 mL of xylene was properly blended before centrifuging. 1.0 mL of the xylene layer was separated, and then the 2,2’-dipyridyl reagent was added and thoroughly mixed. The amount of vitamin E was calculated based on a measurement of absorbance at 460 nm. The reduced glutathione levels were estimated by Boyne and Ellman in 1972 [23]. Briefly, 4.0 mL of metaphosphoric acid were added to 1.0 mL of 10% tissue homogenate. The precipitate that had formed was taken out by centrifuging. 2.0 mL of supernatant received additions of DTNB reagent and disodium hydrogen phosphate, respectively. An absorbance measurement at 412 nm was used to calculate the glutathione content.

The method outlined by Marklund and Marklund in 1974 [24] was used to measure the superoxide dismutase (SOD) activity. Antioxidant enzyme marker (heart) The following components were mixed together: Tris HCl 1.5 mL, pyrogallol 1 mL, and homogenate of heart tissue 1 mL. The absorbance was calculated at 420 nm. The method suggested by Aebi (1984) was employed to evaluate the properties of catalase [25]. In brief, 20 mL of serum and 4 mL of phosphate buffer were combined, and the resulting solution was then incubated at 25 °C. This solution received a 10 mM H2O2 addition, totaling 0.65 mL. The absorbance was calculated at 240 nm. The quantity of GPx in tissue was calculated using the technique described by Rotruck et al. in 1973 [26]. 0.2 mL of Tris buffer, 0.2 mL of EDTA, and 0.1 mL of sodium azide were each added to 0.5 mL of tissue homogenate. This solution was then combined with 0.2 mL of glutathione and 0.1 mL of hydrogen peroxide, and the mixture was incubated at 37 °C for 10 min. Then, 10 minutes of centrifugation at 3000 rpm followed the addition of 0.5 mL of a 10% TCA solution. 2.0 mL of the resulting supernatant was mixed with 1.0 mL of Elman’s (DTNB) reagent and 3.0 mL of disodium dihydrogen phosphate. The absorbance was measured and calculated at 412 nm. The Carlberg and Mannervik approach (1985) [27] was used to assess GR concentration. In a nutshell, 0.5 mL of phosphate buffer, 50 L of GSSG, 50 L of NADPH, and 10 L of distilled water were combined with 10 L of tissue homogenate. The absorbance was measured and calculated at 340 nm. The Habig et al. (1974) technique was used to calculate GST [28]. One millilitre of the sample, two and a half millilitres of phosphate buffer, one and a half millilitres of GSH, and one and a half millilitres of CDNB were added, mixed, and allowed to sit for five minutes. Measurement and computation of absorbance were done.

### 2.11. Measurement of MI markers in the serum

Five minutes were given for the mixture to sit after the addition of one millilitre of the sample, 2.5 millilitres of phosphate buffer, 1.5 millilitres of GSH, and 1.5 millilitres of CDNB. The absorbance was calculated and measured.

### 2.12. Measurement of aldose reductase activity

In order to isolate aortas for further investigation, animals were euthanized with an anaesthetic overdose (200 mg/kg of ketamine mixed with 40 mg/kg of xylazine administered intraperitoneally). Aortic homogenates were subjected to spectrophotometric analysis to determine the aldose reductase activity, as previously discussed [29]. A Bradford assay was used to quantify the protein concentration using bovine serum albumin as a reference.

### 2.13. Measurement of glyoxalase 1 (GLO-1) activity

According to a method previously published by McLellan and Thornalley [30], GLO-1 activity was measured using spectrophotometry. For 10 min at 25°C, the increase in absorbance at 240 nm caused by the production of S-D-lactoylglutathione was observed.

### 2.14. Measurement of methylglyoxal (MG)

According to the previously stated protocols, MG (a precursor in the development of AGE) was detected using HPLC methods in the neutralised perchloric acid extracts of heart tissue. [31].

### 2.15. Measurement of AGE

Cardiac tissues were homogenised in 1 ml of 1×9 PBS after being rinsed with 19 PBS, and they were then kept overnight at 20 C. The homogenates were centrifuged for 5 min. at 50009g and 2-8 °C after two freeze-thaw cycles were used to rupture the cell membranes. A supernatant sample was taken for analysis. AGE levels were measured in homogenates and serum using an ELISA kit designed specifically for use with rats (ABIN368041, antibodies-online GmbH, Aachen, Germany) in accordance with the manufacturer’s instructions.

### 2.16. Measurement of HMGB1

HMGB1 levels were measured using a commercial kit (HMGB1 ELISA kit A76696, Sigma-Aldrich). The assay was performed as described in the manufacturer’s protocol.

### 2.17. Measurement of RAGE

A commercial kit (Rat RAGE/AGER ELISA, #RAB0009, Sigma-Aldrich) was used to quantify RAGE levels. Briefly, concentrated plasma was used, and the test was carried out in accordance with the manufacturer’s protocol.

### 2.18. Measurement of pro-inflammatory cytokines in the serum

TNFα and IL6 serum levels were determined using an ELISA commercial kit in accordance with the manufacturer’s instructions.

### 2.19. Measurement of Heart Mitochondrial Enzymes

The heart’s mitochondrial components were isolated using the adapted methods of Fontana-Ayoub M (2005) [32]. Small pieces of cardiac tissue were mixed with 2 mL of isolation buffer B briefly. The tissue homogenate was combined with 3 mL of isolation buffer B, and the combination was centrifuged at 800 g for approximately 10 min at 4 °C. The supernatant was then separated and centrifuged once more at 10,000 g for roughly 10 min at 4 °C. The supernatant was separated. Isolation buffer A was used to resuspend the mitochondrial suspension in a volume of 500 L. Once more, centrifugation was done at 10,000 g and 4 °C for around 10 min. The mitochondrial suspension was resuspended in 200 L of suspension buffer. For about ten minutes, a second 10,000-g centrifugation at 4 °C was performed. The mitochondrial suspension was reconstituted in 200 L of suspension buffer. In order to later identify the mitochondrial enzymes, a 5 l volume of the mitochondrial suspension was transferred to a 2 mL container and kept at 20 °C. Isocitrate dehydrogenase (IDH) action was studied using the approach described by Bell and Baron (1960) [33]. The following ingredients were added in order: 0.3 mL manganous chloride, 0.2 mL substrate, 0.2 mL mitochondrial suspension, and 0.4 mL Tris-HCl buffer. This mixture was then incubated for 60 min followed by the addition of 1.0 mL of DNPH and then 0.5 mL of EDTA solution. After 20 min,10.0 mL of 0.4 NaOH was added. At 390 nm, the color’s intensity was measured. The Reed and Mukherjee (1969) technique was utilized to measure the enzyme’s ketoglutarate dehydrogenase activity [34]. Briefly, magnesium sulfate, thiamine pyrophosphate, potassium ferricyanide, and -ketoglutarate were combined with 0.15 mL of phosphate buffer. This solution was mixed with 0.2 mL of mitochondrial suspension, and it was then incubated at 30 °C for 30 min. Before centrifuging, 1 mL of 10% TCA was further added. 0.1 millilitres of potassium ferricyanide, 0.5 millilitres of ferric ammonium sulphate-dupanol reagent, and 1 millilitre of 4% dupanol were added after the supernatant had been separated.

### 2.20. Measurement of Tissue Collagen Content

After combining 0.1 ml of EDTA, 0.1 ml of bovine albumin, 1 ml of phosphate buffer, 0.2 ml of potassium ferricyanide, 0.3 ml of sodium succinate, and 0.1 ml of potassium cyanide, 0.2 ml of mitochondrial suspension was added. Colour intensity was measured and calculated at 420 nm. Malate dehydrogenase’s role was investigated by Mehler et al. in 1948 [35]. 0.3 mL of buffer, 0.1 mL of oxaloacetate, and 0.1 mL of NADH were mixed, and then 0.1 mL of mitochondrial suspension was added. Colour intensity was measured and calculated at 340 nm. A 20% solution in methyl cellosolve that had been warmed to 60 °C to help in solubilization was added, thoroughly mixed, and then put in a water bath for 20 min at 60 °C after chilling in tap water for 5 min. The acquired colour at 557 nm was measured spectrophotometrically in comparison to a distilled water blank. Standard solutions were treated identically to produce a calibration curve. The collagen content of the tissue samples was determined and represented as mg/gm tissue by multiplying the hydroxyproline concentration by 8 [36].

### 2.21. Histopathological Study

According to protocol, heart tissue was prepared for histopathology slides and photographs were taken. The hearts and lipids were carefully extracted after euthanasia. They were immediately fixed for 48 hours in 10% buffered formalin, gradually dried by immersion in progressively higher and higher concentrations of water-ethanol, cleaned with xylene, and then once again embedded in paraffin before being cut into 5–6 m slices with a microtome. The sections were stained using, among other things, hematoxylin and eosin dyes [37,38].

### 2.22. Statistical analysis

The results of the analysis were presented as mean±SD (Standard Deviation). Dunnett’s test was used to compare all groups to the control group after one-way ANOVA (Graph Pad Instat, USA) statistical analysis.

## 3. Result

### 3.1. Heart:Body Weight Ratio

A mathematically calculated value in which the heart’s total weight is divided by the total body weight, and the result is shown as a ratio which is termed as the heart: body weight ratio. The heart:body weight ratio was statistically highly significant (p < 0.001) in isoproterenol group rats (ISG) when compared to (NCG). The rosuvastatin-treated group showed a statistically very significant (p < 0.01) decrease in heart:body weight ratio when compared to the isoproterenol group (ISG). The per se group (PSG) showed no significant (p > 0.05) change in terms of H:B ratio as compared with the normal control group (NCG) (Figure 1).

**Figure 1.**
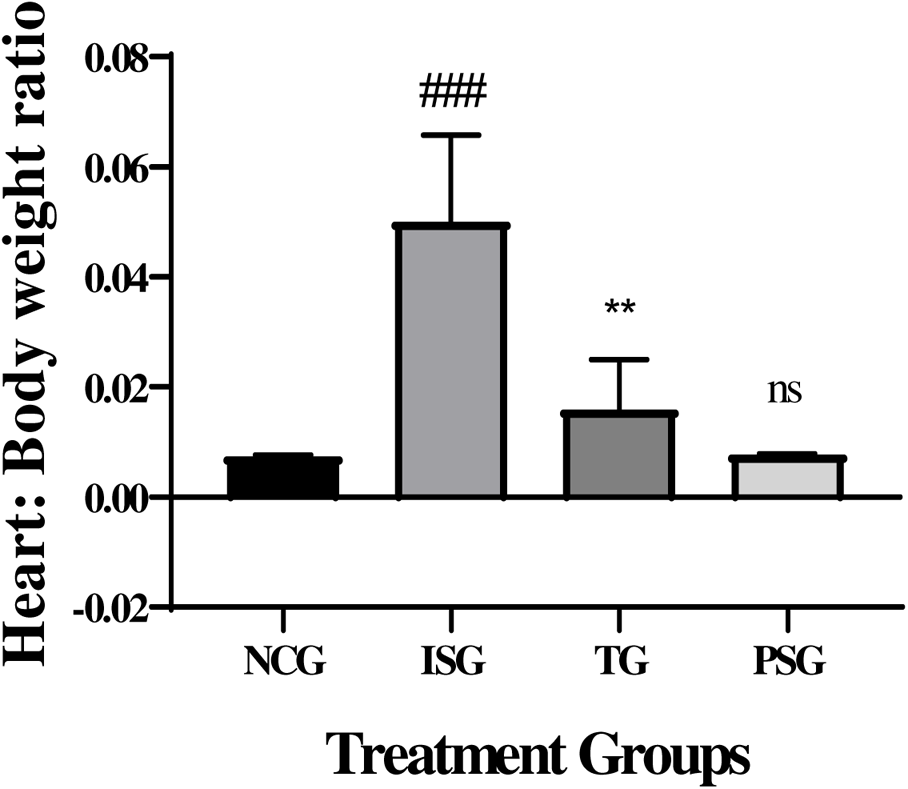
Heart: Body weight ratio in the different groups. All values are expressed as mean±SD; (n=6) in each group. Data was subjected to one-way ANOVA followed by Dunnett’s test when normal control group (NCG) was compared to isoproterenol control group (ISG), treatment group (TG) was compared to isoproterenol control group (ISG), while per se (PSG) was compared to normal control group (NCG). ^ns^p>0.05, *p<0.05, **p<0.01, ***p<0.001 when compared to normal control group (NCG), *p<0.05, **p<0.01, ***p<0.001 when compared to isoproterenol control group (ISG).

### 3.2. Echocardiography measurement

Isoproterenol induced hearts showed markedly damaged at the end of the study. As expected, LVEF and FS were worse in the iso groups than in the normal control group, demonstrating systolic dysfunction (Table 1). In the same way, we saw a progressive increase in LV diameters over the time in ISG group. Isoproterenol induced hearts had infarcted area which was decreased in treatment group. After 28 days of treatment with rosuvastatin the echocardiography markers were found to be near-normal levels. The rosuvastatin-treaded group (10 mg/kg) significantly reversed those results. The PSG group showed no significant difference when compared to the NCG group.

**Table 1.** Ejection fraction, Fractional shortening, LV systolic diameter, LV diastolic diameter, Infarct size of treatment group.

|  | NCG | ISG | TG | PSG |
| --- | --- | --- | --- | --- |
| Ejection fraction (%) | 64.6 | 32.3<sup>###</sup> | 49.1<sup>***</sup> | 63.9<sup>ns</sup> |
| Fractional shortening (%) | 37.2 | 19.7<sup>###</sup> | 31.5<sup>***</sup> | 38.6<sup>ns</sup> |
| LV systolic diameter (mm) | 3.7 | 7.9<sup>###</sup> | 6.8<sup>***</sup> | 3.9<sup>ns</sup> |
| LV diastolic diameter (mm) | 7.8 | 12.7<sup>###</sup> | 11.4<sup>***</sup> | 12.4<sup>ns</sup> |
| Infarct size (%) | — | 42±1.7<sup>###</sup> | 12±0.8<sup>***</sup> | — |

All values are expressed as mean±SD; (n=6) in each group. Data was subjected to one-way ANOVA followed by Dunnett’s test when normal control group (NCG) was compared to isoproterenol control group (ISG), treatment group (TG) was compared to isoproterenol control group (ISG), while per se (PSG) was compared to normal control group (NCG). ^ns^p>0.05, *p<0.05, **p<0.01, ***p<0.001 when compared to normal control group (NCG), *p<0.05, **p<0.01, ***p<0.001 when compared to isoproterenol control group (ISG).

### 3.3. Serum Biochemical Markers

The activities of serum biochemical markers AST, ALT LDH, and CK-MB was found to increase in the ISG group when compared with NCG group rats. After 28 days of treatment with rosuvastatin the biochemical markers were found to be declined to near-normal levels. The outcome indicates clearly that the rosuvastatin prevented the myocardial tissue from enzymatic leakage from the cell sites, evidencing its protective effect on the myocardium. PSG group showed no significant differences (p > 0.05) in the level of AST, ALT, LDH, and CK-MB when compared to the NCG group (Figure 2).

**Figure 2.**
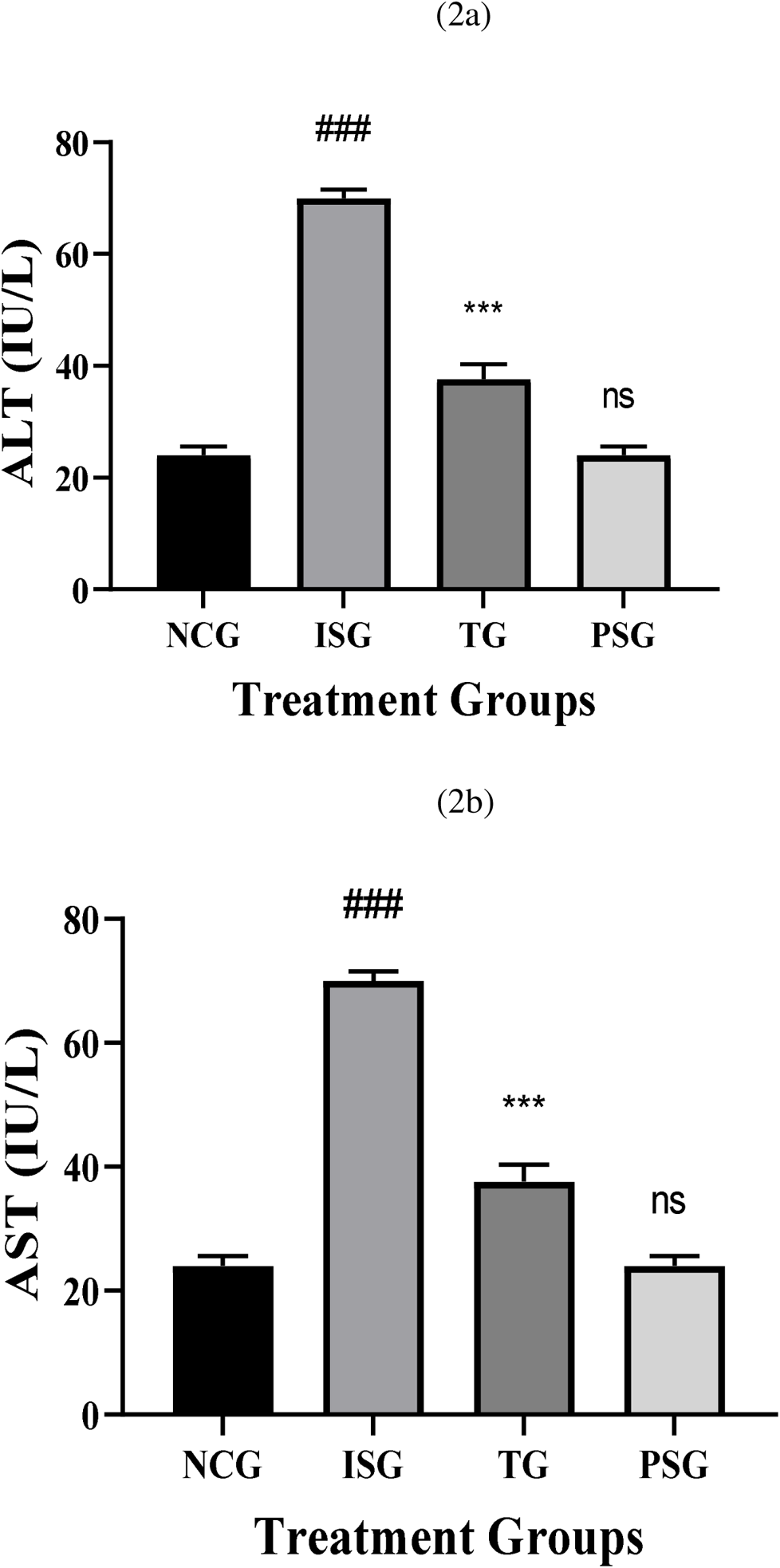
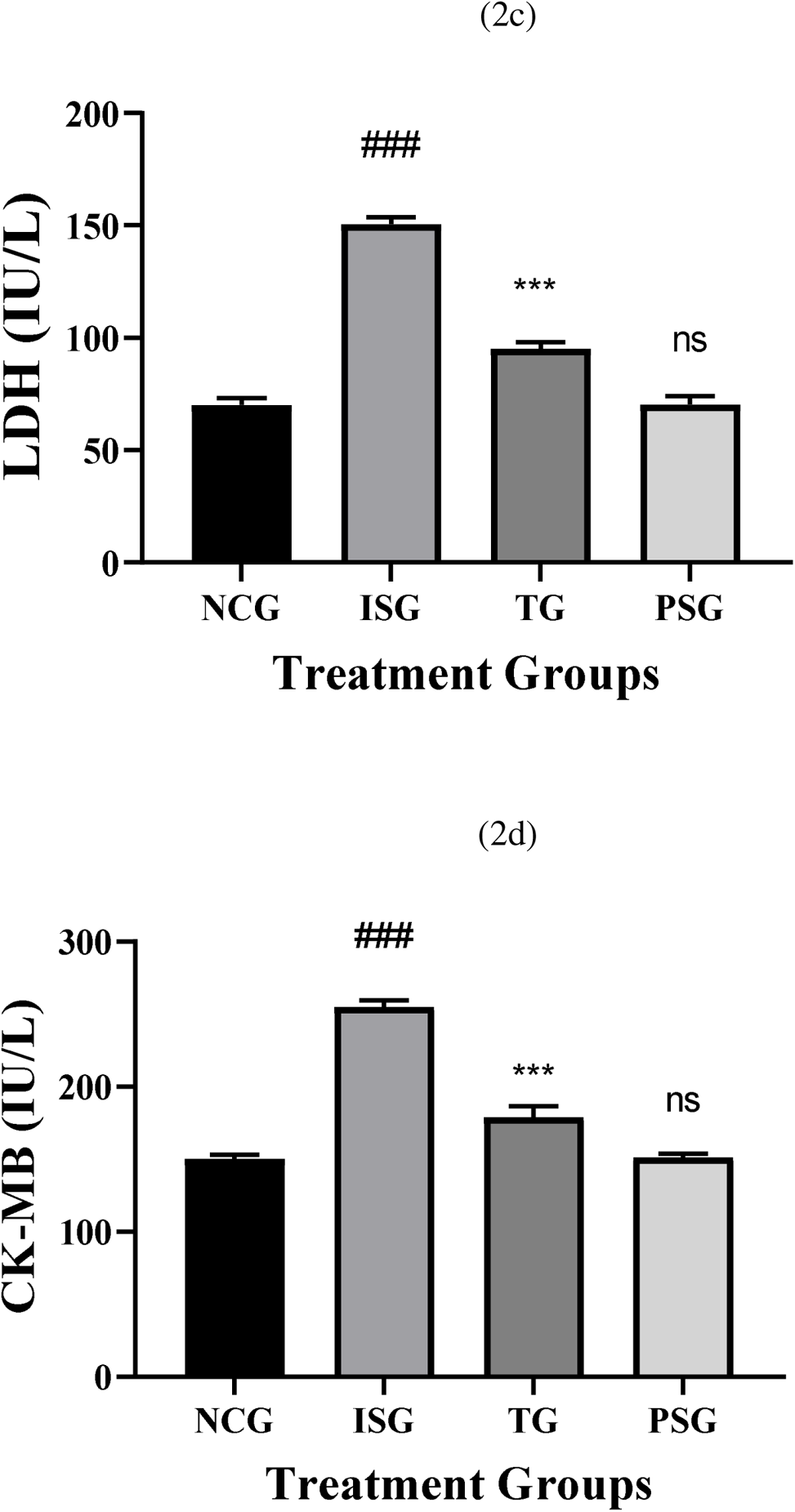
Serum Biochemical estimations in the different groups. All values are expressed as mean±SD; (n=6) in each group. Data were subjected to one-way ANOVA followed by Dunnett’s test when normal control group (NCG) was compared to isoproterenol control group (ISG), treatment group (TG) was compared to isoproterenol control group (ISG), while per se (PSG) was compared to normal control group (NCG). ^ns^p>0.05, *p<0.05, **p<0.01, ***p<0.001 when compared to normal control group (NCG), *p<0.05, **p<0.01, ***p<0.001 when compared to isoproterenol control group (ISG).

### 3.4. Oxidative Stress Parameters

A significant upsurge in the levels of lipid peroxidative marker in serum (TBARS, lipid hydroperoxides, conjugated dienes) in ISG group rats when compared to NCG was observed due to accumulation of free radicals. Rosuvastatin group showed decreased levels of lipid peroxidative marker. The PSG group showed no significant difference in the level of lipid peroxidative marker when compared to the NCG group. The results indicate the high free radical scavenging property or strong neutralizing effect of sericin (Figures 3).

**Figure 3.**
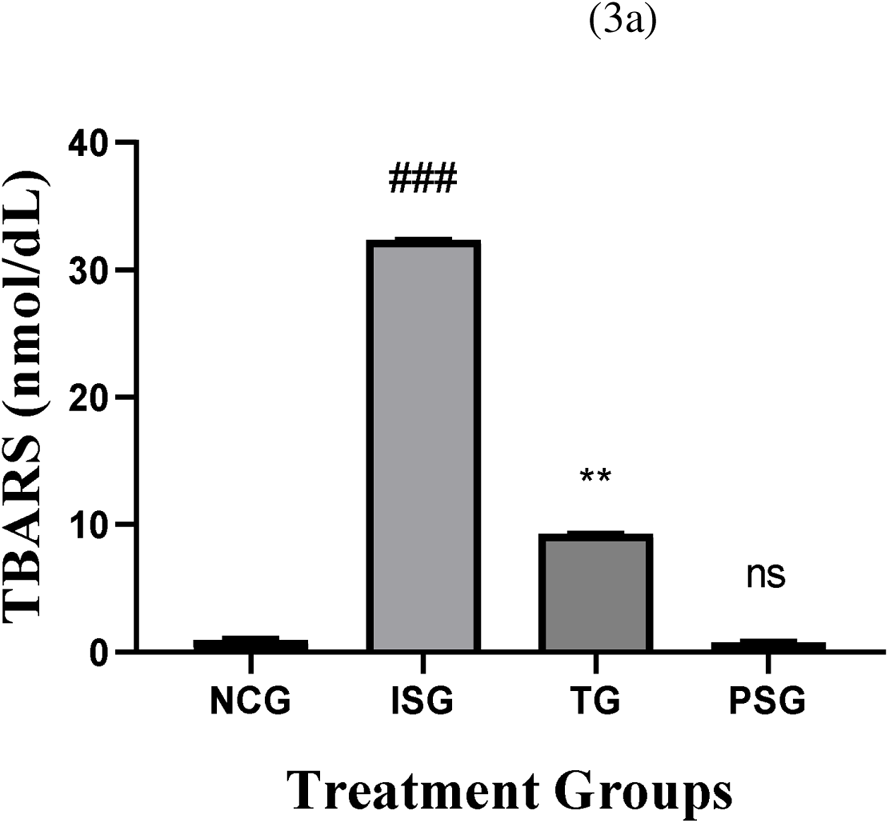
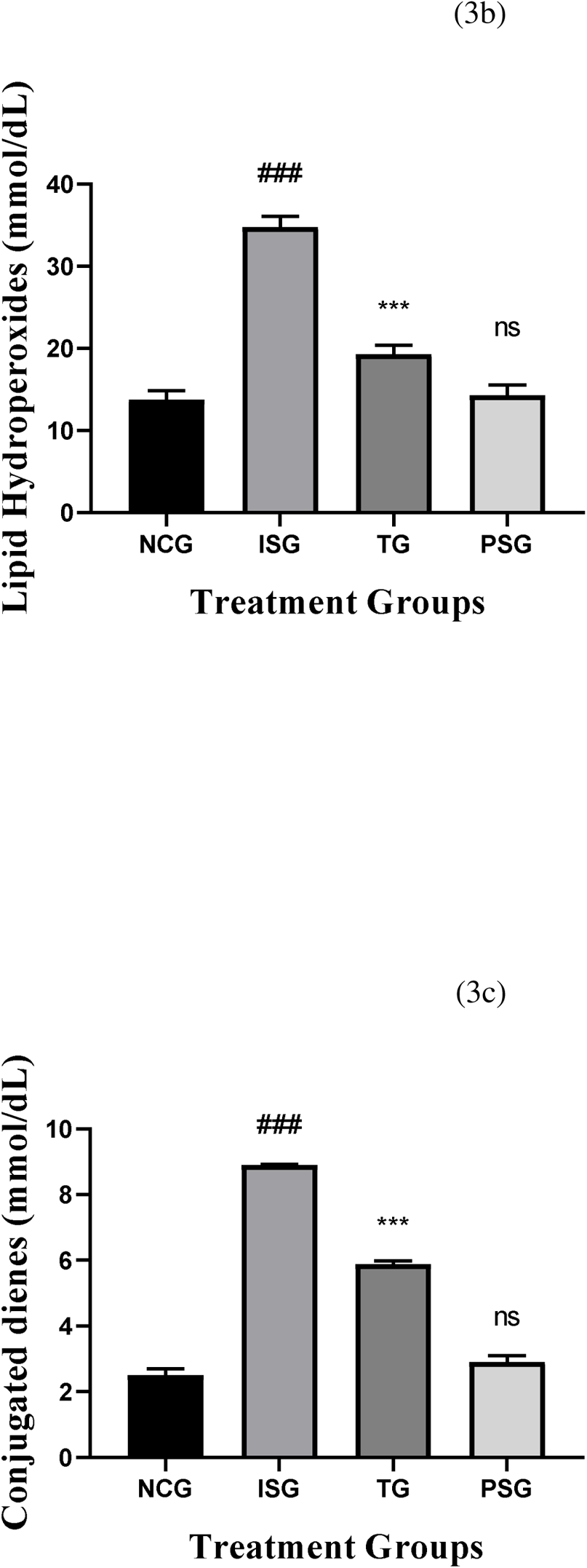
Lipid peroxide markers in serum in the different groups. All values are expressed as mean±SD; (n=6) in each group. Data were subjected to one-way ANOVA followed by Dunnett’s test when normal control group (NCG) was compared to isoproterenol control group (ISG), treatment group (TG) was compared to isoproterenol control group (ISG), while per se (PSG) was compared to normal control group (NCG). ^ns^p>0.05, *p<0.05, **p<0.01, ***p<0.001 when compared to normal control group (NCG), *p<0.05, **p<0.01, ***p<0.001 when compared to isoproterenol control group (ISG).

### 3.5. Oxidative Stress Parameters

AGE accelerates oxidative stress by interacting with a specific receptor RAGE on vascular cells. A significant upsurge in the levels of lipid peroxidative marker in cardiac tissue (TBARS, lipid hydroperoxides, conjugated dienes) in ISG group rats when compared to NCG was observed due to accumulation of free radicals were observed. Treatment group showed decreased levels of lipid peroxidative marker. The PSG group showed no significant difference in the level of lipid peroxidative marker when compared to the NCG group. The results indicate the high free radical scavenging property or strong neutralizing effect of rosuvastatin (Figures 4).

**Figure 4.**
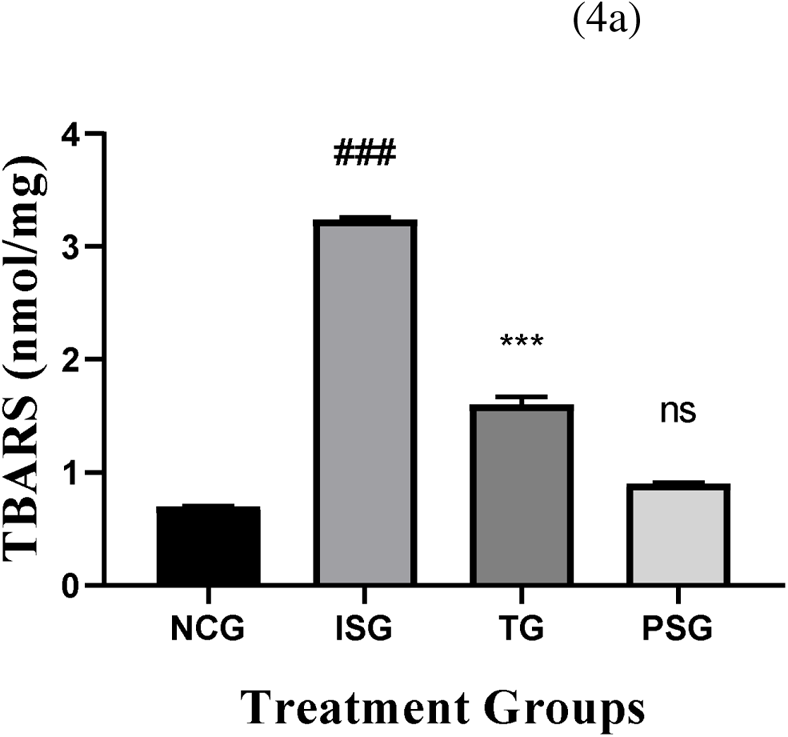
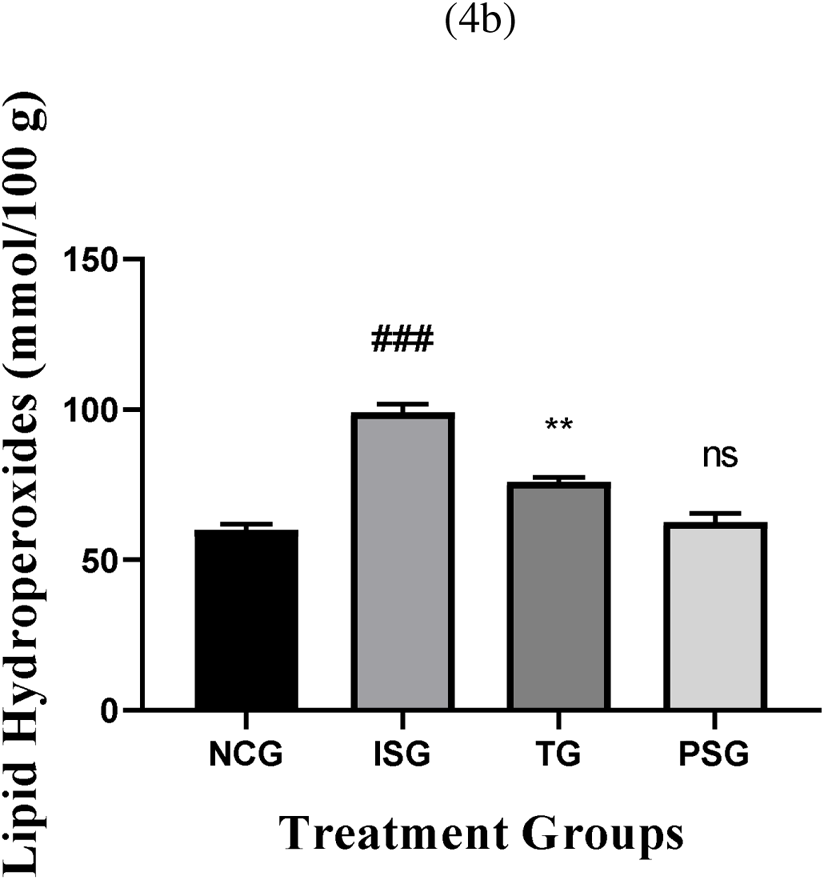
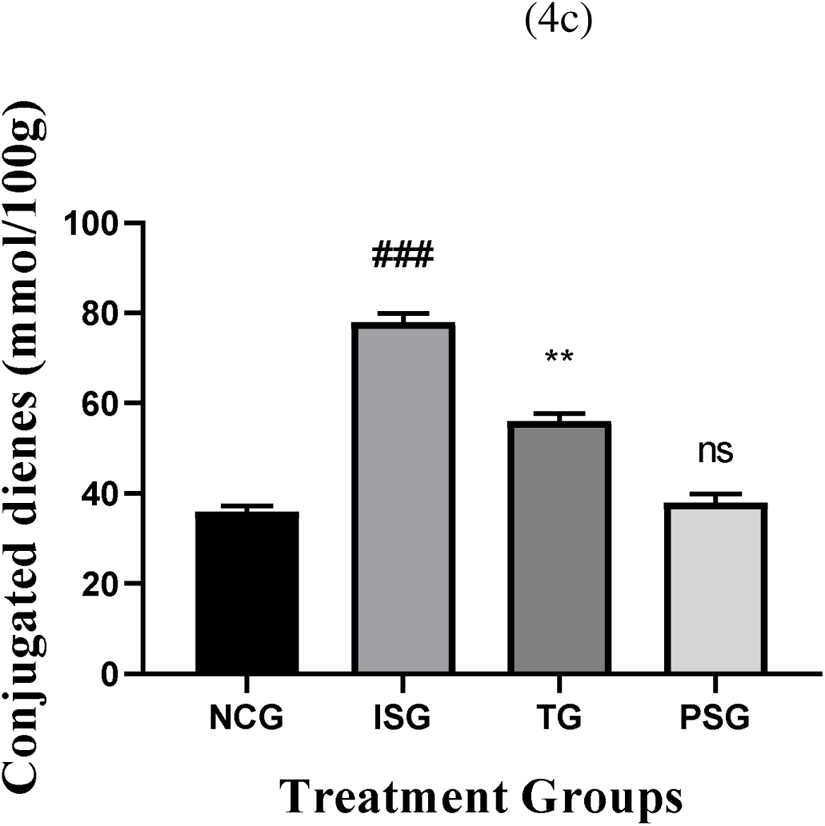
Lipid peroxide markers in tissue in the different groups. All values are expressed as mean±SD; (n=6) in each group. Data were subjected to one-way ANOVA followed by Dunnett’s test when normal control group (NCG) was compared to isoproterenol control group (ISG), treatment group (TG) was compared to isoproterenol control group (ISG), while per se (PSG) was compared to normal control group (NCG). ^ns^p>0.05, *p<0.05, **p<0.01, ***p<0.001 when compared to normal control group (NCG), *p<0.05, **p<0.01, ***p<0.001 when compared to isoproterenol control group (ISG).

### 3.6. Antioxidant Parameters

#### 3.6.1. Non-Enzymatic Antioxidants Marker in Serum

The levels of non-enzymatic antioxidants (viz. vitamin C, vitamin E, and glutathione) were found to be decreased in the serum and myocardial tissue of groups intoxicated with ISO group when compared with the NCG group. The PSG group did not show any changes in antioxidant levels when compared to the NCG group. Treatment with rosuvastatin reversed the total amount of non-enzymatic antioxidants to near-normal levels in the serum/plasma and myocardial tissue thus proving that rosuvastatin possesses good anti-oxidant potential against the action of AGE-RAGE (Figure 5 and 6).

**Figure 5.**
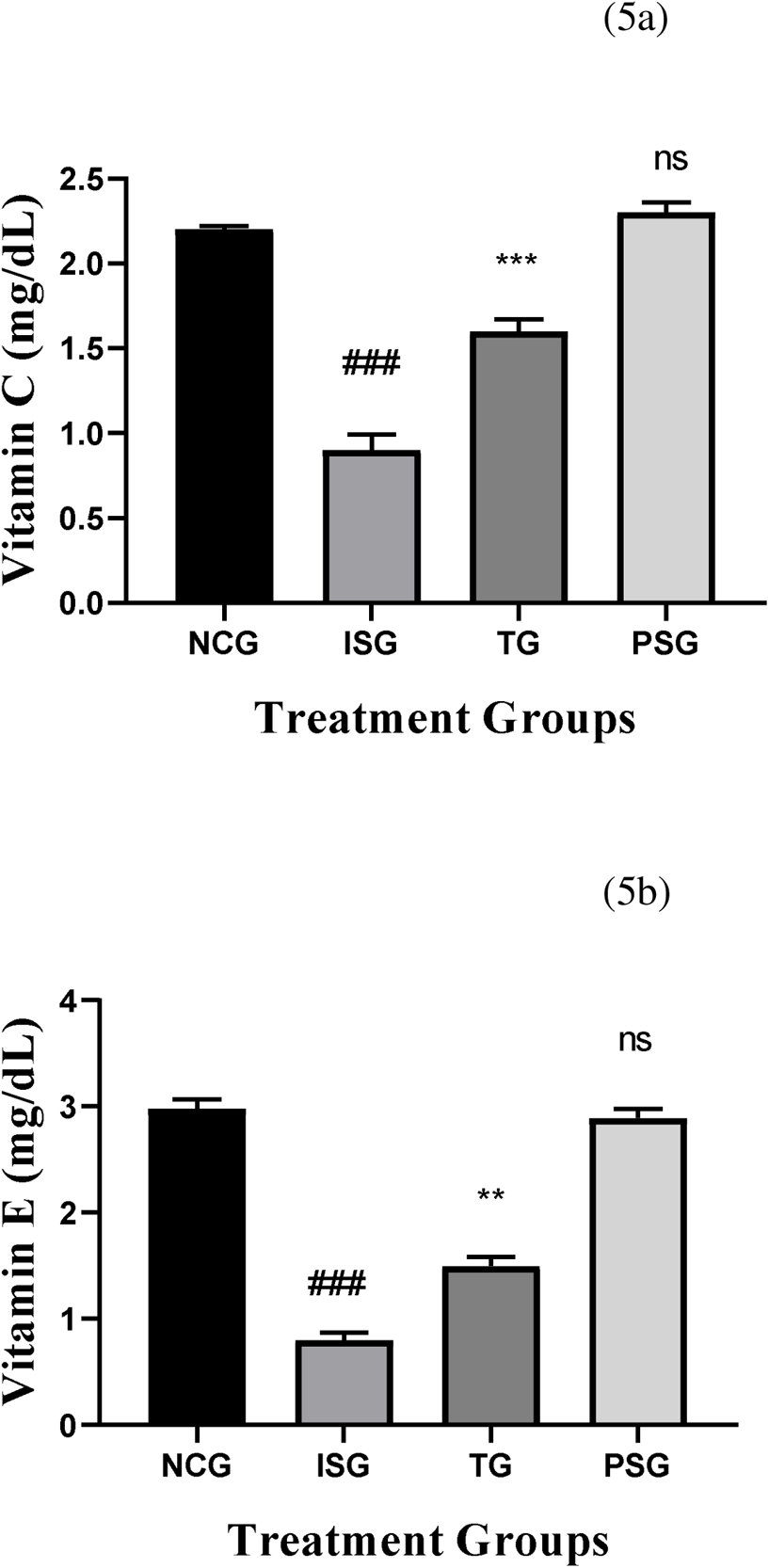
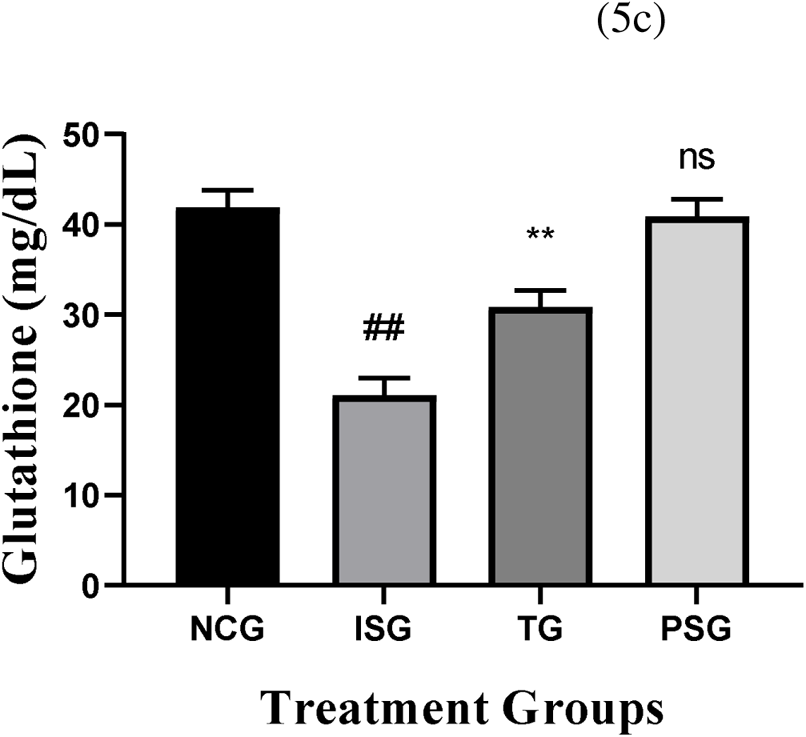
Non-Enzymatic Antioxidant Marker in serum in the different groups. All values are expressed as mean±SD; (n=6) in each group. Data were subjected to one-way ANOVA followed by Dunnett’s test when normal control group (NCG) was compared to isoproterenol control group (ISG), treatment group (TG) was compared to isoproterenol control group (ISG), while per se (PSG) was compared to normal control group (NCG). ^ns^p>0.05, *p<0.05, **p<0.01, ***p<0.001 when compared to normal control group (NCG), *p<0.05, **p<0.01, ***p<0.001 when compared to isoproterenol control group (ISG).

**Figure 6.**
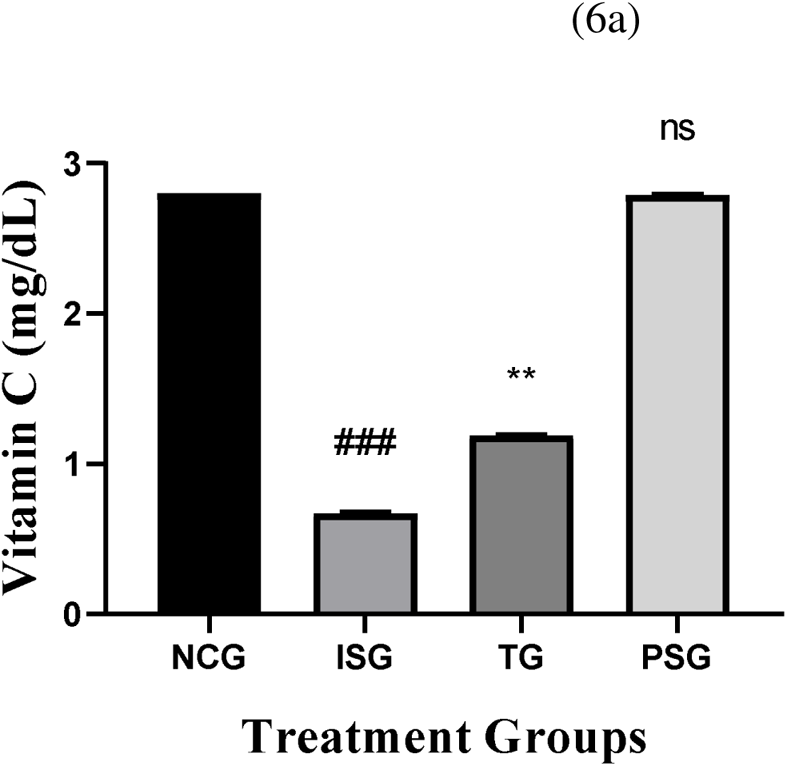
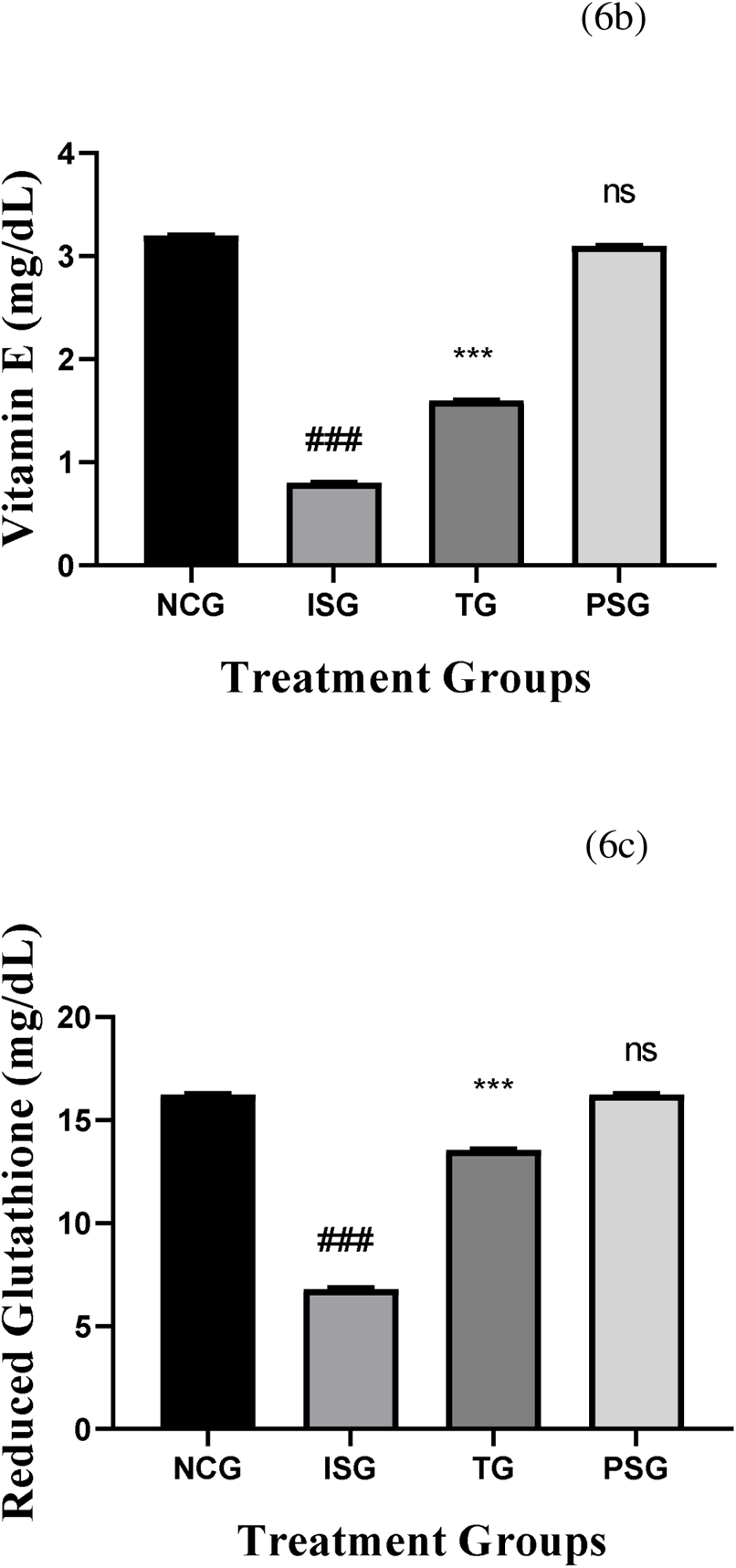
Non-Enzymatic Antioxidant Marker in heart tissue in the different groups. All values are expressed as mean±SD; (n=6) in each group. Data were subjected to one-way ANOVA followed by Dunnett’s test when normal control group (NCG) was compared to isoproterenol control group (ISG), treatment group (TG) was compared to isoproterenol control group (ISG), while per se (PSG) was compared to normal control group (NCG). ^ns^p>0.05, *p<0.05, **p<0.01, ***p<0.001 when compared to normal control group (NCG), *p<0.05, **p<0.01, ***p<0.001 when compared to isoproterenol control group (ISG).

#### 3.6.2. Antioxidant Marker (Enzymatic) in Heart

The number of enzymatic antioxidants such as (GR), (CAT), (GPx), (SOD), and (GST) were decreased in ISG group when compared with NCG rats. Treatment with rosuvastatin for a period of 28 days restored the actions of these enzymes to near-normal levels when compared to ISG group rats. The PSG group did not show any changes in antioxidant levels when compared to the NCG group (Figure 7).

**Figure 7.**
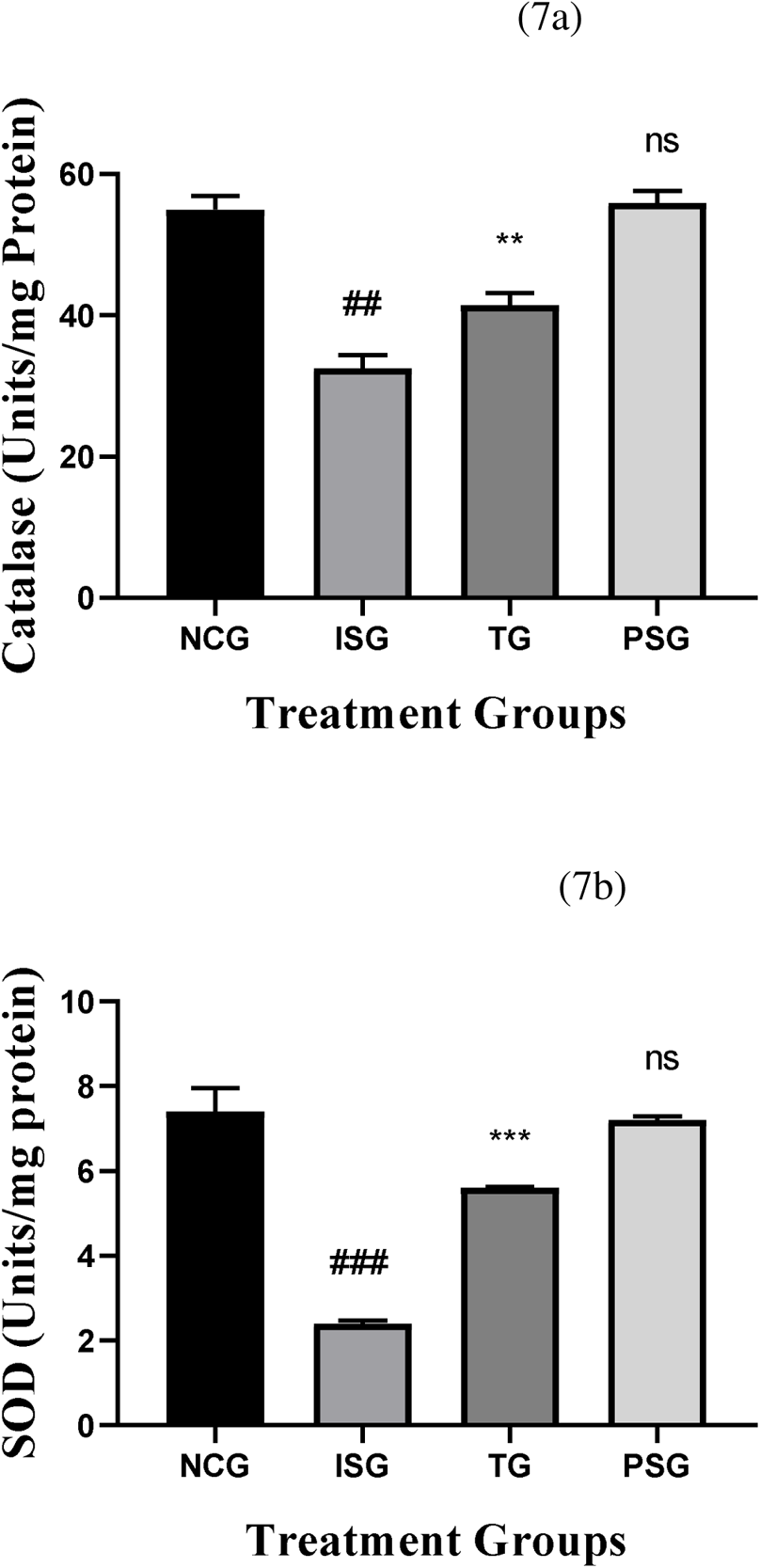
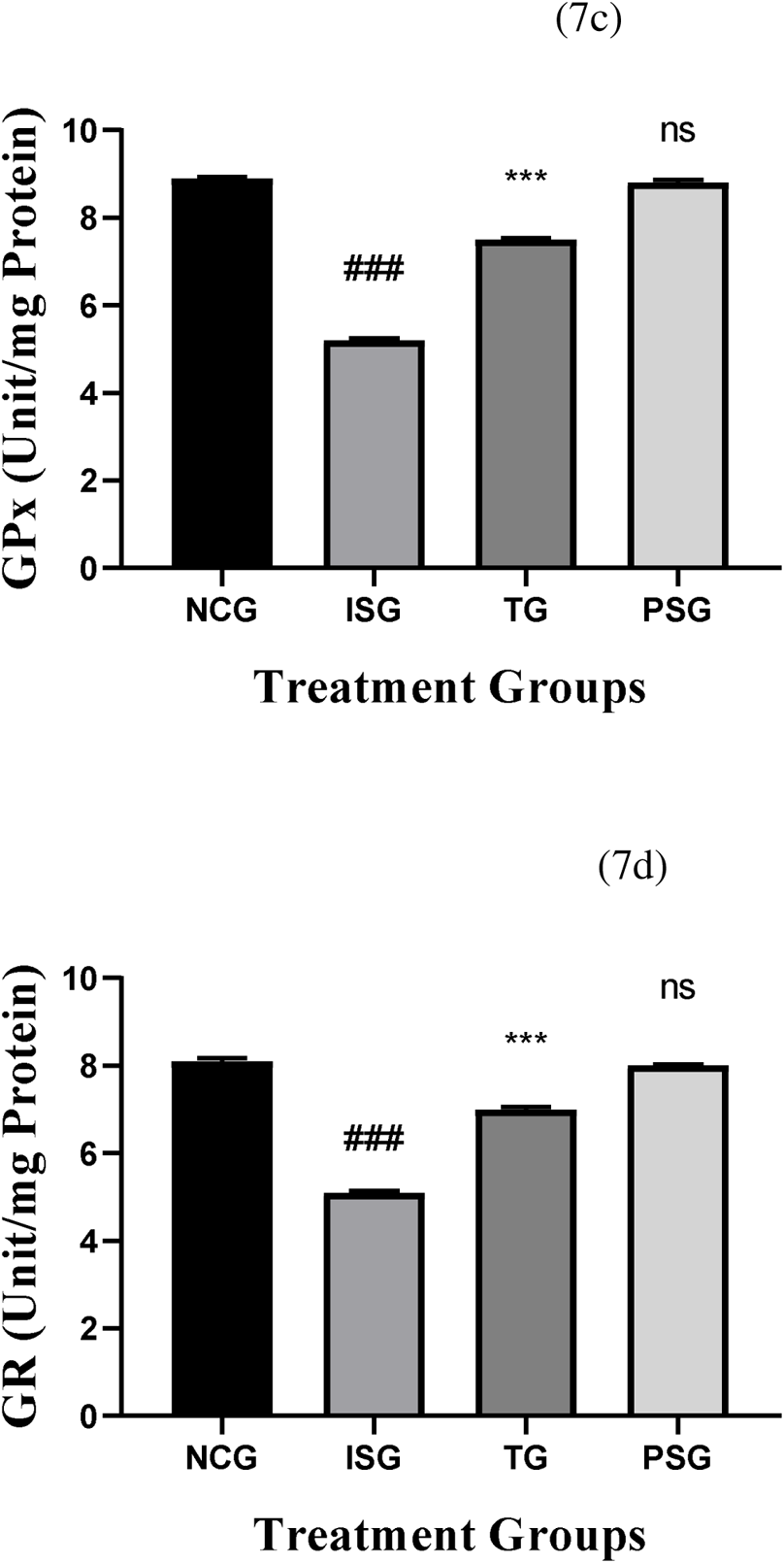
Enzymatic Antioxidant Marker in heart tissue in the different groups. All values are expressed as mean±SD; (n=6) in each group. Data were subjected to one-way ANOVA followed by Dunnett’s test when normal control group (NCG) was compared to isoproterenol control group (ISG), treatment group (TG) was compared to isoproterenol control group (ISG), while per se (PSG) was compared to normal control group (NCG). ^ns^p>0.05, *p<0.05, **p<0.01, ***p<0.001 when compared to normal control group (NCG), *p<0.05, **p<0.01, ***p<0.001 when compared to isoproterenol control group (ISG).

### 3.7. Measurement of AGE

The serum ISG group exhibited enhanced activity of aldose reductase, reduced activity of Glo-1, increased levels of MG and AGE (Fig. 1d), and upregulated serum RAGE and, which indicated that AGE–RAGE pathway was activated in ISG group. The possible presence of AGEs in the serum samples of the animal model for glycation was verified with ELISA kit. The presence of the serum AGEs was observed to be highly significant in ISG induced group when compared with NCG group. Treatment with rosuvastatin ameliorated the serum AGE level when compared with iso group. The PSG group did not show any changes in AGE levels when compared to the NCG group (Figure 8).

**Figure 8.**
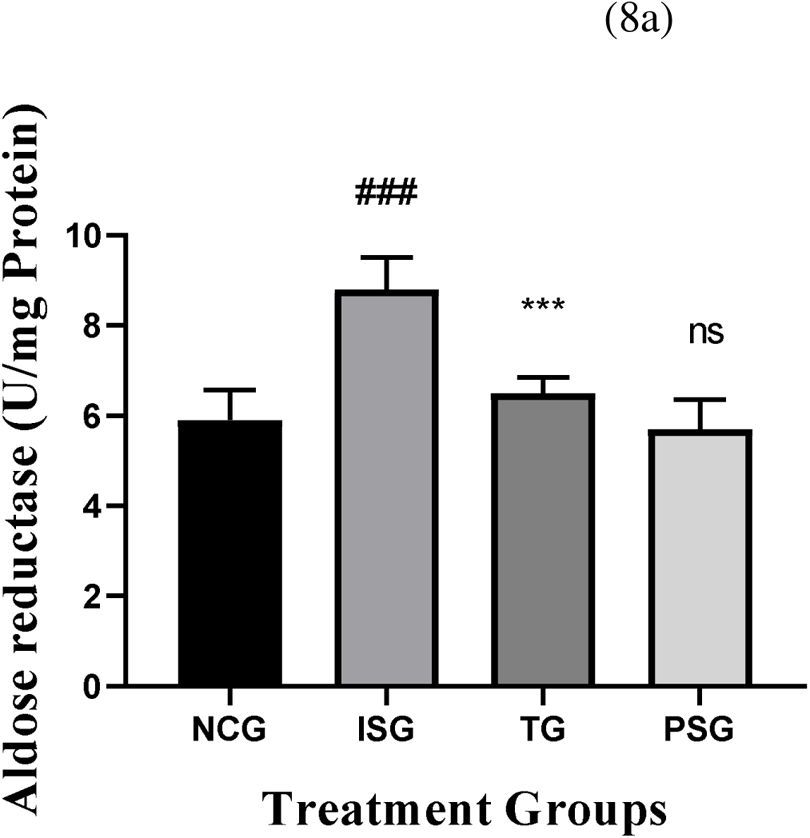
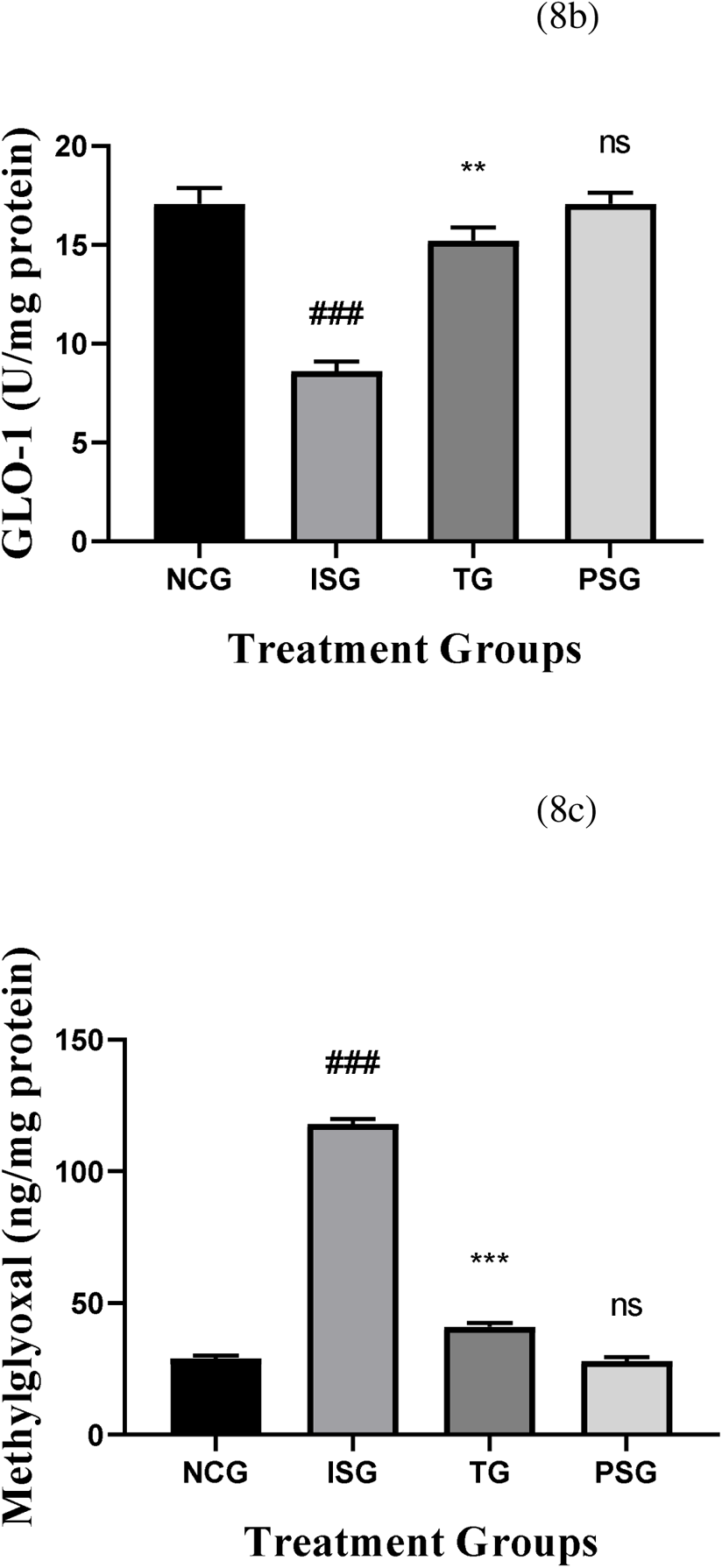
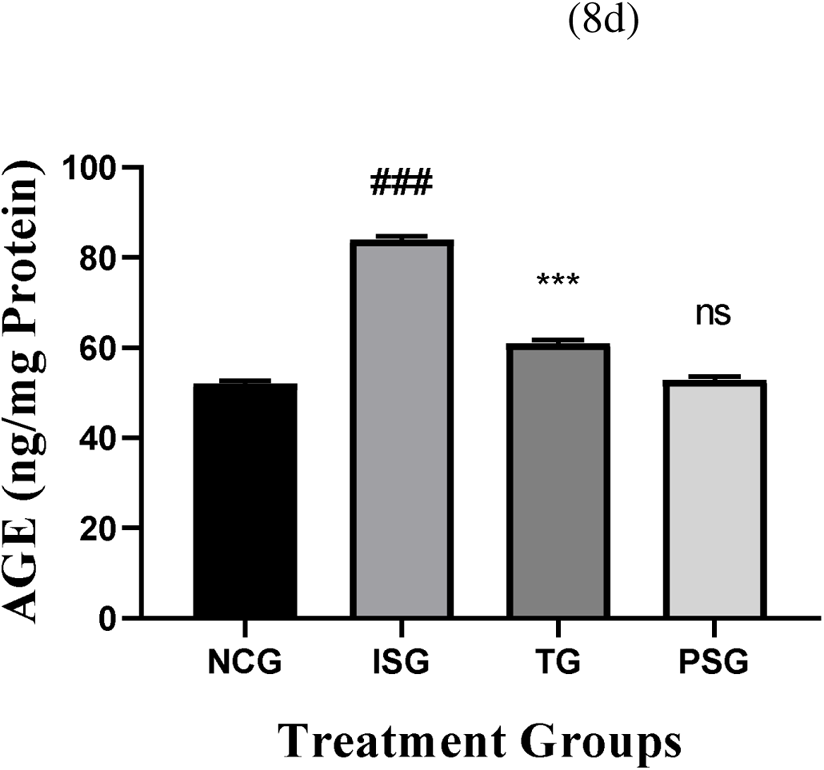
Measurement of AGE pathway in the different groups. All values are expressed as mean±SD; (n=6) in each group. Data were subjected to one-way ANOVA followed by Dunnett’s test when normal control group (NCG) was compared to isoproterenol control group (ISG), treatment group (TG) was compared to isoproterenol control group (ISG), while per se (PSG) was compared to normal control group (NCG). ^ns^p>0.05, *p<0.05, **p<0.01, ***p<0.001 when compared to normal control group (NCG), *p<0.05, **p<0.01, ***p<0.001 when compared to isoproterenol control group (ISG).

### 3.8. Measurement of RAGE

To further explore the role of rosuvastatin in protecting cell death, we measured the expression of its target genes. We initially focused on the RAGE gene. Total RAGE levels in serum increased approximately 50% in the iso group when compared with the NCG group. Treatment with rosuvastatin ameliorated the serum RAGE level when compared with iso group. The PSG group did not show any changes in RAGE levels when compared to the NCG group (Figure 9).

**Figure 9.**
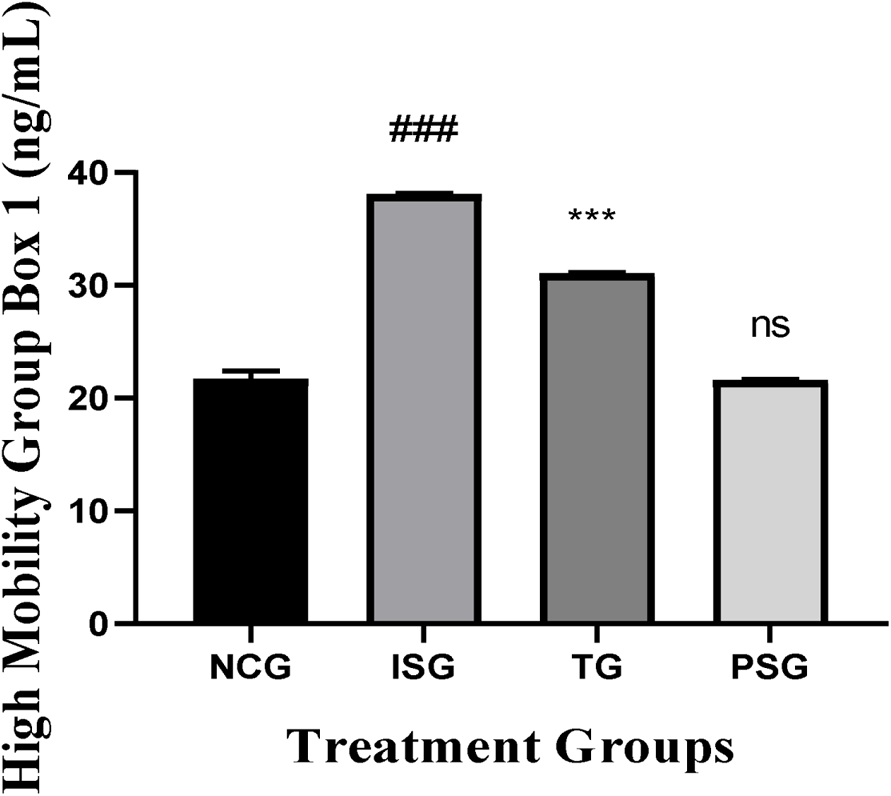
High mobility group box 1 in the different groups. All values are expressed as mean±SD; (n=6) in each group. Data were subjected to one-way ANOVA followed by Dunnett’s test when normal control group (NCG) was compared to isoproterenol control group (ISG), treatment group (TG) was compared to isoproterenol control group (ISG), while per se (PSG) was compared to normal control group (NCG). ^ns^p>0.05, *p<0.05, **p<0.01, ***p<0.001 when compared to normal control group (NCG), *p<0.05, **p<0.01, ***p<0.001 when compared to isoproterenol control group (ISG).

### 3.9. Measurement of HMGB1

High-mobility group box-1 (HMGB1) is a ligand for the receptor for advanced glycation endproducts (RAGE). An HMGB1-RAGE interaction has been implicated in cardiac dysfunction. Compared to the normal control group, administration of Isoproterenol significantly increased the HMGB1 levels in the heart. Pretreatment with rosuvastatin decreased ISO-induced elevation of serum HMGB1. The PSG group did not show any changes in HMGB1 levels when compared to the NCG group (Figure 10).

**Figure 10.**
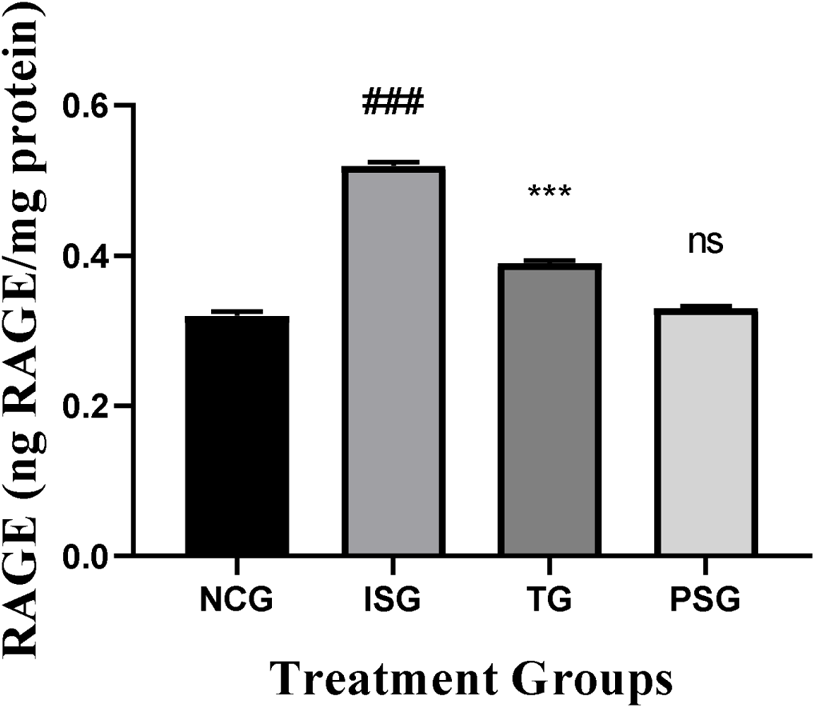
RAGE levels in the different groups. All values are expressed as mean±SD; (n=6) in each group. Data were subjected to one-way ANOVA followed by Dunnett’s test when normal control group (NCG) was compared to isoproterenol control group (ISG), treatment group (TG) was compared to isoproterenol control group (ISG), while per se (PSG) was compared to normal control group (NCG). ^ns^p>0.05, *p<0.05, **p<0.01, ***p<0.001 when compared to normal control group (NCG), *p<0.05, **p<0.01, ***p<0.001 when compared to isoproterenol control group (ISG).

### 3.10. Effect of rosuvastatin on cytokines in heart

Compared to the normal control group, administration of ISO significantly increased the secretion levels of TNF-α and IL-6 in the heart. Pretreatment with rosuvastatin decreased ISO-induced elevation of cardiac TNF-α. The PSG group did not show any changes in cytokines levels when compared to the NCG group (Figure 11).

**Figure 11.**
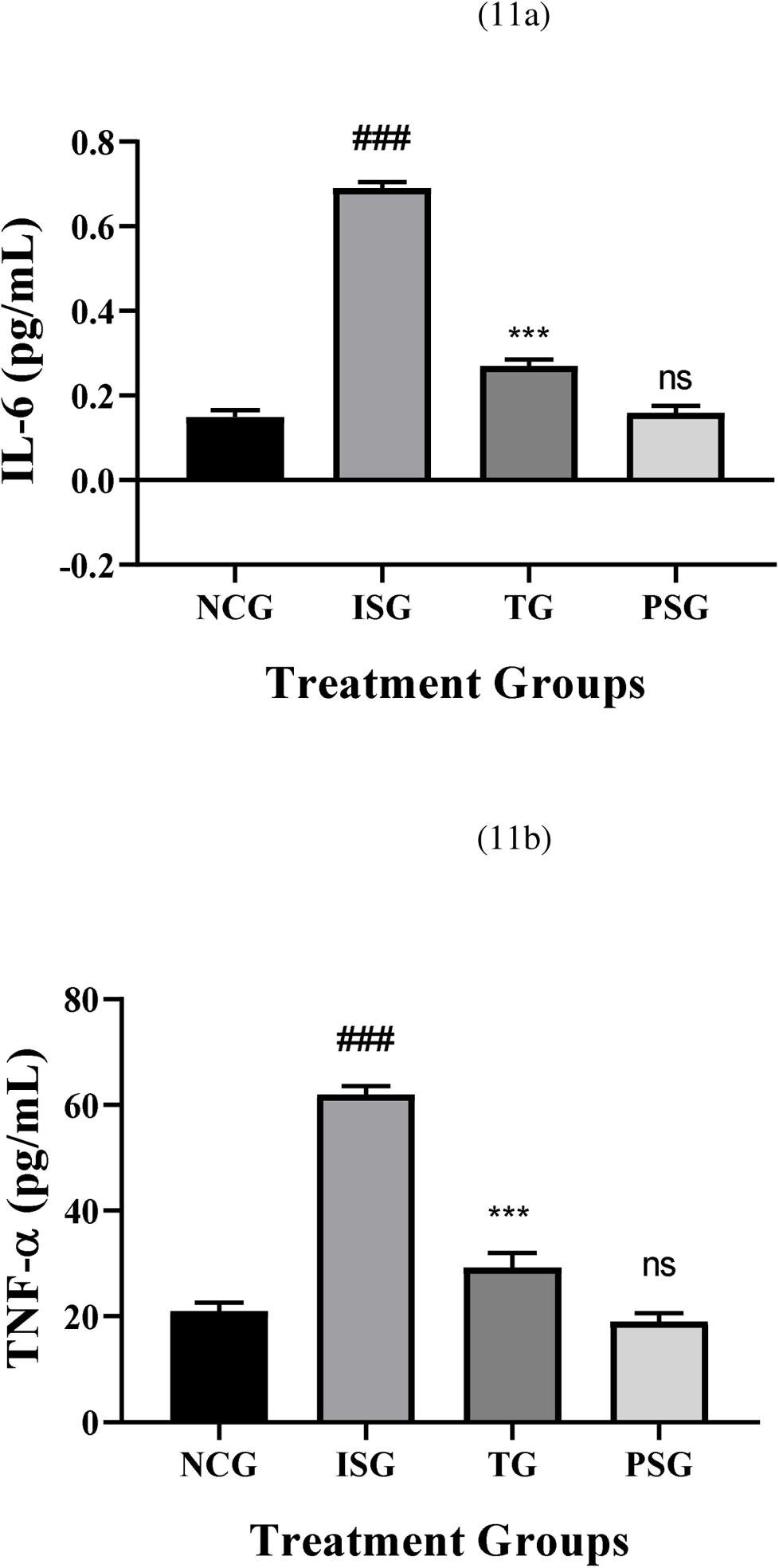
Pro-inflammatory cytokines in the serum in the different groups. All values are expressed as mean±SD; (n=6) in each group. Data were subjected to one-way ANOVA followed by Dunnett’s test when normal control group (NCG) was compared to isoproterenol control group (ISG), treatment group (TG) was compared to isoproterenol control group (ISG), while per se (PSG) was compared to normal control group (NCG). ^ns^p>0.05, *p<0.05, **p<0.01, ***p<0.001 when compared to normal control group (NCG), *p<0.05, **p<0.01, ***p<0.001 when compared to isoproterenol control group (ISG).

### 3.11. Total Myocardial Collagen Content and Fibrosis Estimation

Total myocardial collagen content and fibrosis were estimated and were found to be statistically highly significant in isoproterenol group rats (ISG). The treatment group showed a statistically very significant decrease in collagen content and fibrosis. No significant changes in the level of collagen content and fibrosis were observed in the PSG group. The result suggests that rosuvastatin reduces the synthesis of collagen in myocardiocytes, thus reducing the incidence of fibrosis, reduction of fibrosis and synthesis of collagens contributes to the protective effect of rosuvastatin against hypertrophy (Figure 12).

**Figure 12.**
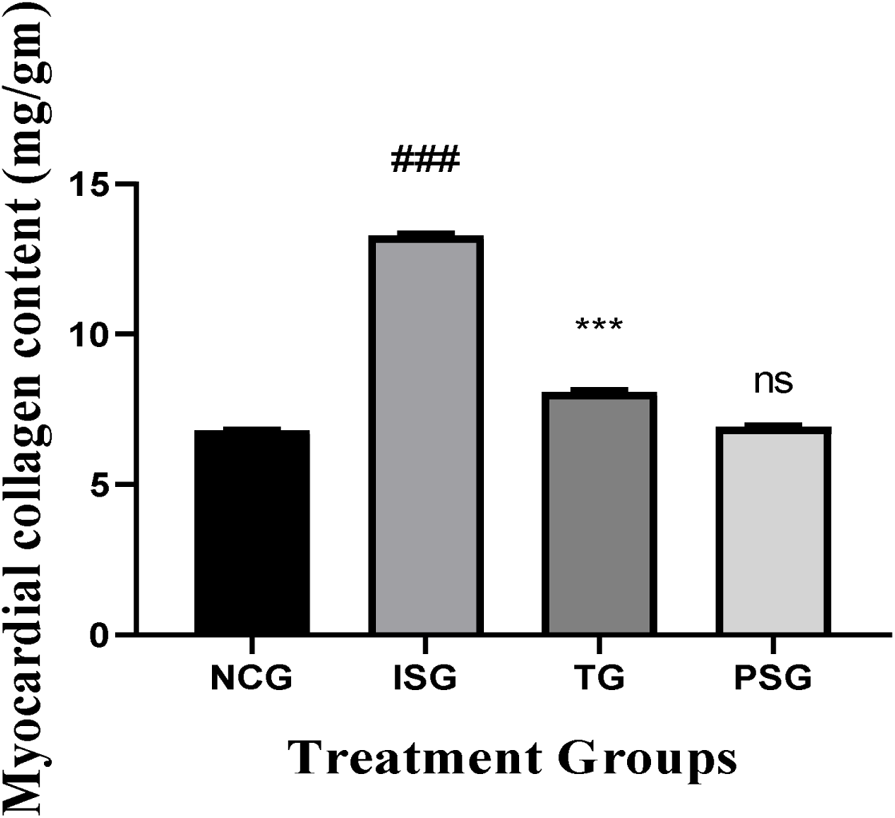
Myocardial collagen content in the different groups. All values are expressed as mean±SD; (n=6) in each group. Data were subjected to one-way ANOVA followed by Dunnett’s test when normal control group (NCG) was compared to isoproterenol control group (ISG), treatment group (TG) was compared to isoproterenol control group (ISG), while per se (PSG) was compared to normal control group (NCG). ^ns^p>0.05, *p<0.05, **p<0.01, ***p<0.001 when compared to normal control group (NCG), *p<0.05, **p<0.01, ***p<0.001 when compared to isoproterenol control group (ISG).

### 3.12. Heart Mitochondrial Enzymes

Various different biochemical enzymes exist in mitochondria, for instance, α-Ketoglutarate dehydrogenase (KDH), Isocitrate dehydrogenase (IDH), Succinate dehydrogenase (SDH), and Malate dehydrogenase (MDH) which were measured in the treated heart group. Mitochondrial enzymes present in the heart were estimated and the isoproterenol group (ISG) showed statistically highly significant (p < 0.001) reduction in the level of MDH, KDH, IDH, and SDH when compared with the normal control group (NCG. In treatment group the level of MDH, KDH, and SDH were statistically very significantly (p < 0.01) while the level of IDH was statistically significantly (p < 0.05) increased when compared with the isoproterenol group (ISG). In the per se group (PSG) exposure of rats to rosuvastatin produced alterations (p > 0.05) in the level of tissue MDH, IDH, KDH, and SDH when compared with animals of the normal control group (NCG) (Figure 13).

**Figure 13.**
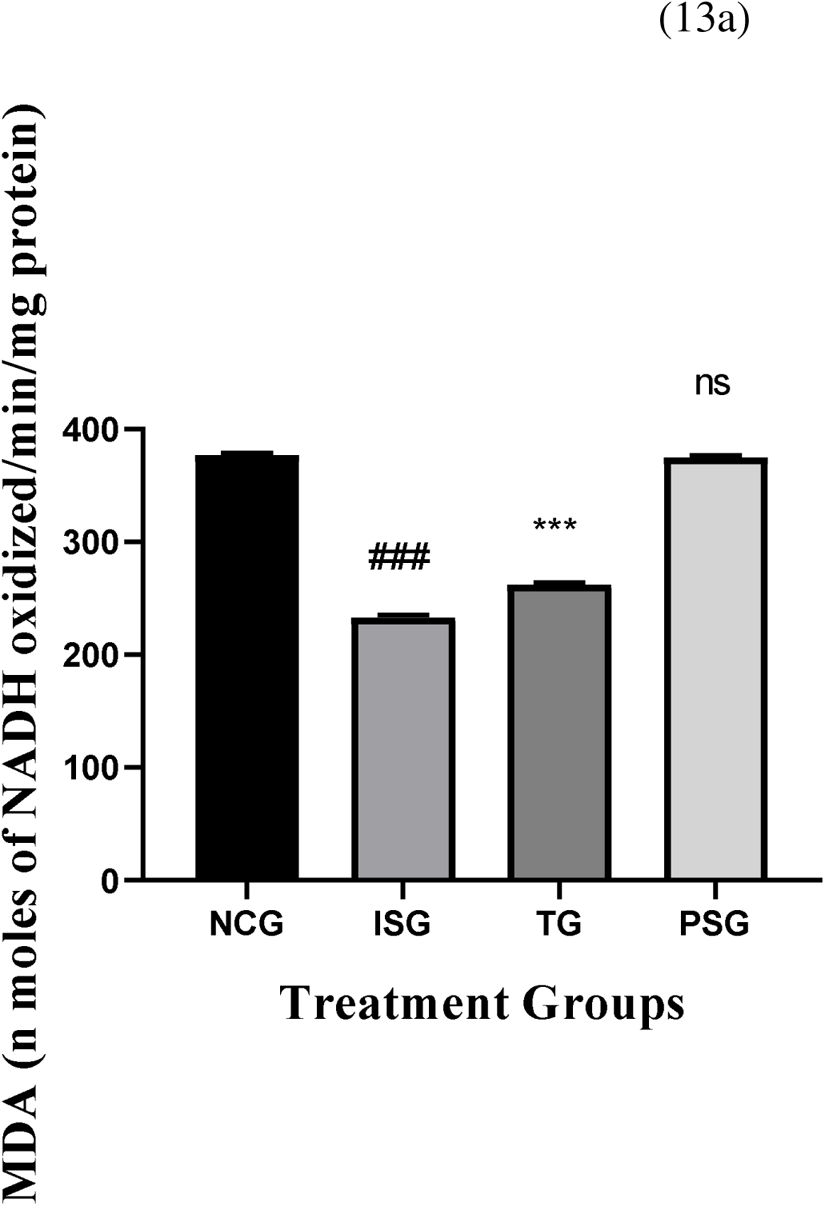
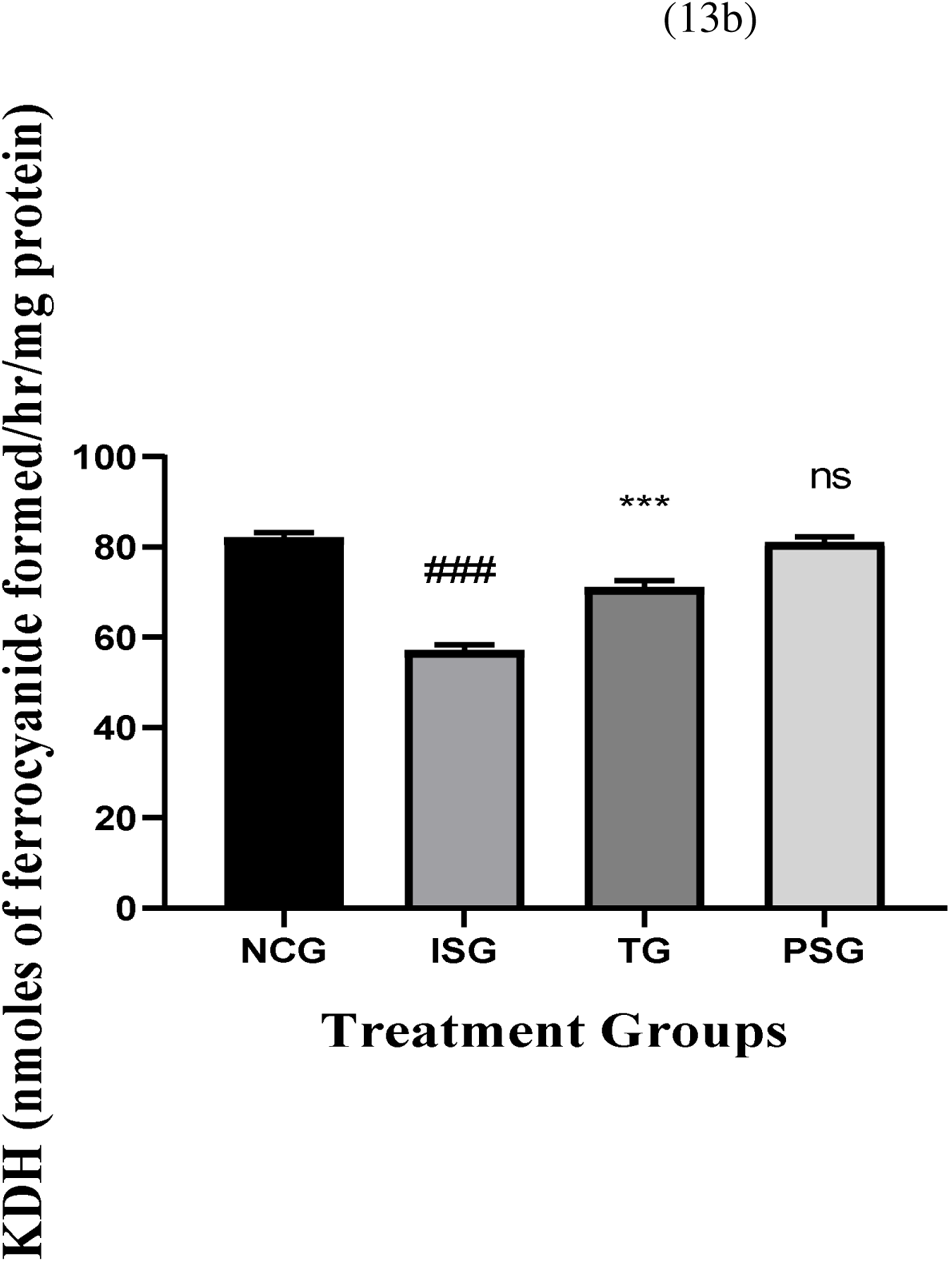
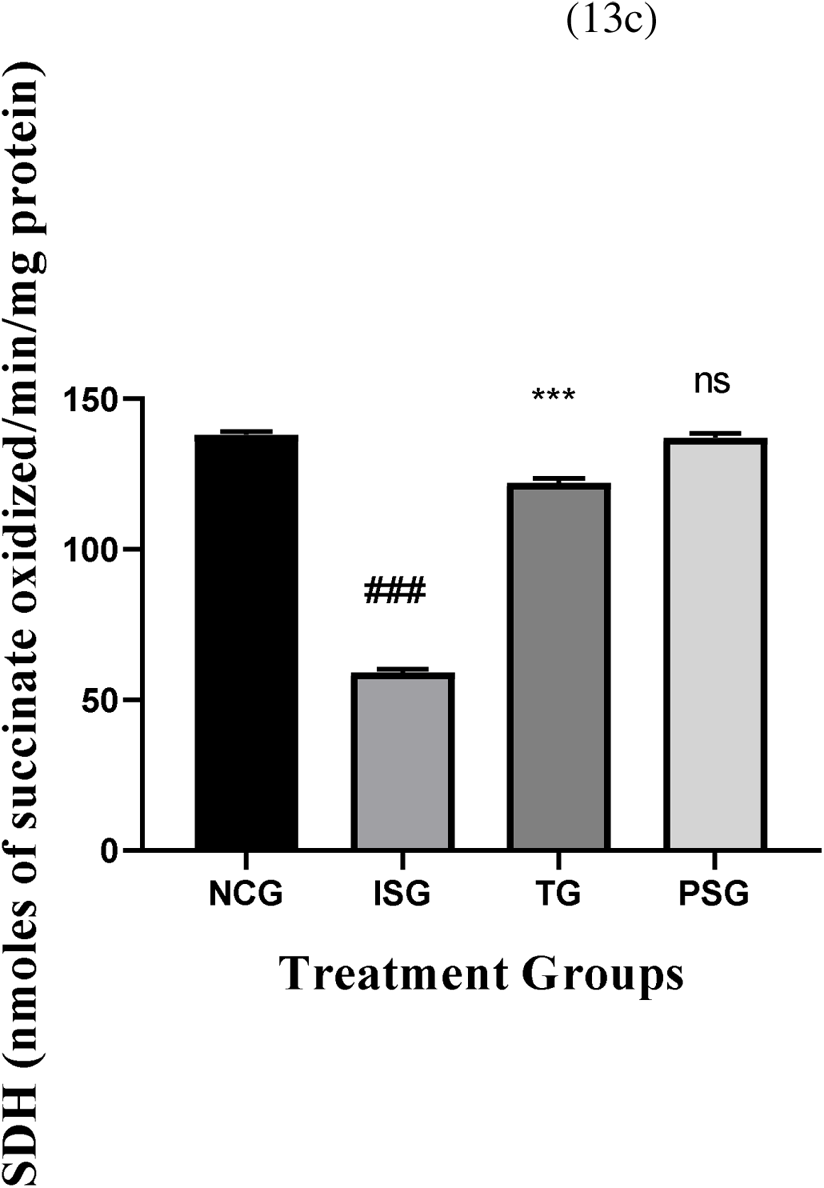
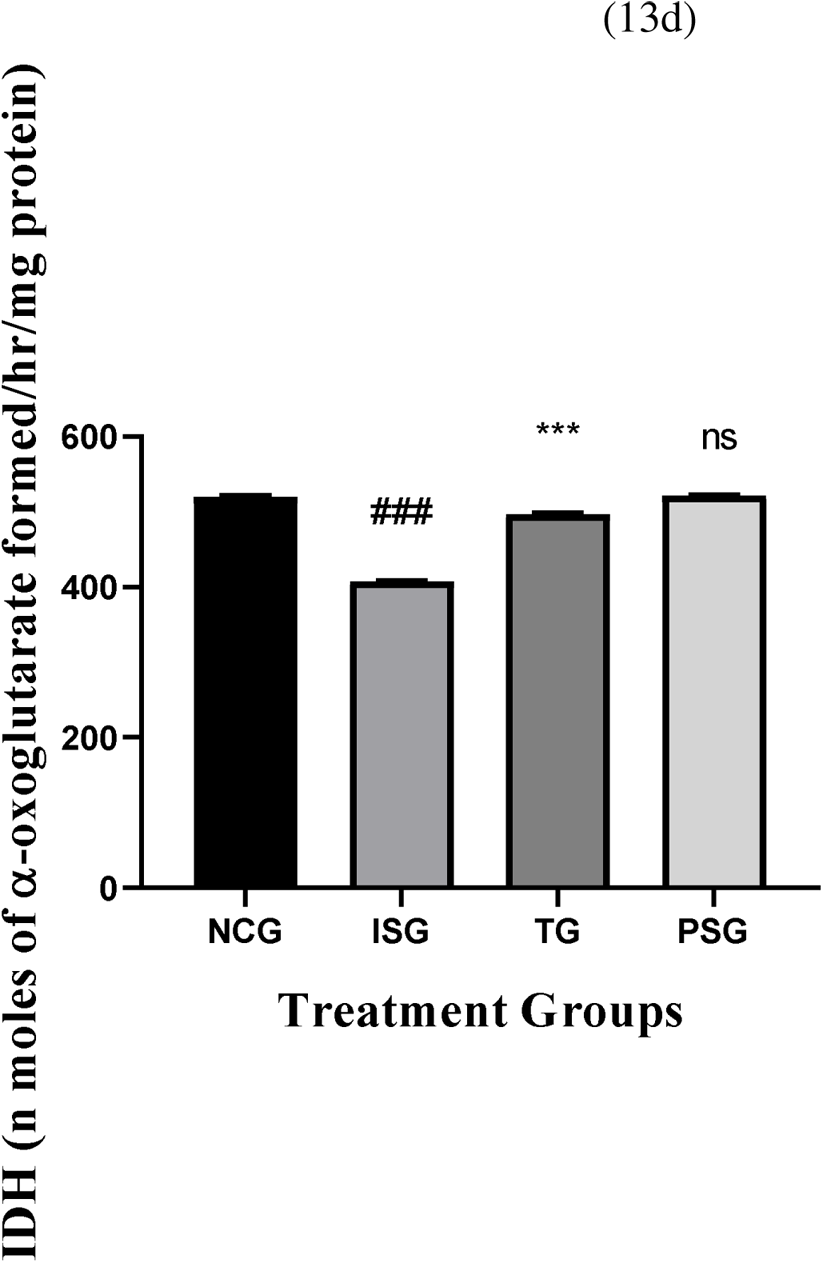
Mitochondrial enzyme levels (heart) in the different groups. All values are expressed as mean±SD; (n=6) in each group. Data were subjected to one-way ANOVA followed by Dunnett’s test when normal control group (NCG) was compared to isoproterenol control group (ISG), treatment group (TG) was compared to isoproterenol control group (ISG), while per se (PSG) was compared to normal control group (NCG). ^ns^p>0.05, *p<0.05, **p<0.01, ***p<0.001 when compared to normal control group (NCG), *p<0.05, **p<0.01, ***p<0.001 when compared to isoproterenol control group (ISG).

### 3.13. Histopathology

The histopathological end result showed that normal control rats administered every day with saline have confirmed the regular alignment of myocardium, endocardium, and epicardium in addition to papillary muscle fibres and vasculature in comparison to isoproterenol (ISG) group which displayed massive infarction with wall lesions thrombi and random acute aneurysms. Whilst the rosuvastatin group confirmed focal lesions without an indication of myonecrosis, myophagocytosis, and lymphocytic infiltration. The per se (PSG) showed regular myocardial tissue structure, a systemized pattern showing no vacuolation and myofiber striae with nuclei in center (Figure 14).

**Figure 14.**
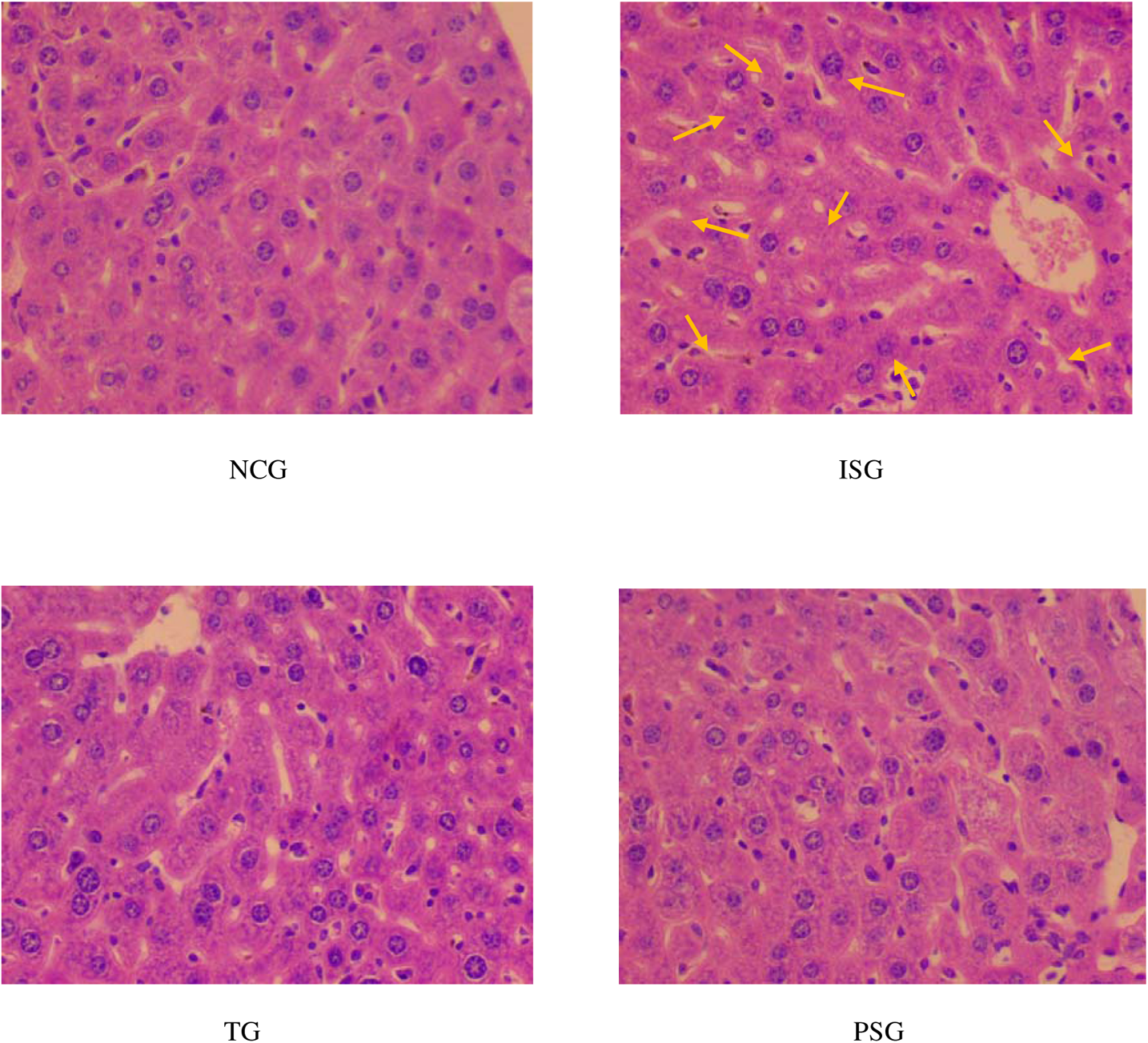
Histopathological images of the heart in different treatment groups. Arrows in ISG shows necrosis and infiltration.

## 4. Discussion

The activation of the AGE/RAGE pathway has a profound impact on the aetiology of cardiovascular disease. In a number of animal models of cardiovascular diseases, reducing AGE-RAGE activation has been decisively demonstrated to be beneficial.

Rosuvastatin, one of the most potent statins, has anti-inflammatory and endothelium protective properties in addition to its well-known lipid-lowering benefits [39–41].

Additionally, there is mounting proof that rosuvastatin can reduce blood AGEs levels and RAGE expression in heart injury [42–44]. This result is thought to be a brand-new molecular mechanism for the pleiotropic effects of rosuvastatin.

Accordingly, the results of the current investigation demonstrated that rosuvastatin treatment enhanced AGE-RAGE, biochemical indices, and histology. These findings are corroborated by earlier research; according to Fei et al. [45], activation of the AGE/RAGE also causes the formation of intracellular oxidative stress, which in turn causes the redox-sensitive transcription factor NF-kappaB to become active. However, the results of this investigation imply that Rosuvastatin exerts its therapeutic effects by focusing on the AGE-RAGE axis. In addition, Kim et al [46] showed that rosuvastatin prevented the long-term detrimental effects of ADR on left ventricular function. Ke et al [47] and Du et al [48] showed that pre- and post-conditioning with rosuvastatin reduces ischemia/reperfusion myocardial injury through the inhibition of HMGB1. In addition, the main effect of statins is to lower the blood cholesterol levels, whereas their pleiotropic effects on AGE-RAGE axis are secondary effects. Isoproterenol-induced myocardial injury has been widely used to investigate the effect of drugs on AGE induced myocardial infraction [49,50]. The current study successfully establisheda rat model of myocardial damage, as shown by drastically elevated blood levels of AST, ALT, CK-MB, and LDH as well as aberrant cardiac microstructure found on histological examination. These outcomes are consistent with earlier in vivo investigations [51]. Rosuvastatin can reduce ISO-induced myocardial infarction injury by blocking the AGE-RAGE axis, which was further supported by the present study’s significantly reduced blood levels of myocardial injury markers and visibly reduced histological changes. Rosuvastatin (10 mg/kg) therapy resulted in a decrease in the blood levels of lactate dehydrogenase and creatine kinase-MB isoenzyme, indicating that rosuvastatin mitigated the severity of myocardial injury and hence restricted the leakage of these enzymes from the myocardium. Rosuvastatins are anticipated to have significant impacts in addition to their potential to decrease cholesterol. The enzyme that limits the rate at which HMG-CoA is converted to mevalonate, 3-hydroxy-3-methylglutaryl-coenzyme-A (HMG-CoA) reductase, is competitively inhibited by rosuvastatins [52]. As a result, they inhibit the synthesis of cholesterol, disrupt the mevalonate pathway, and disrupt the production of numerous other crucial molecules that are necessary for maintaining cellular activities [53].

Oxidative stress and myocardial damage are interlinked. Patients with acute myocardial infarction have been found to have metabolic disturbances in oxidants and antioxidants[54]. We noticed more malondialdehyde and less antioxidants in the rat cardiac tissues that underwent ISO treatment as a result of AGE-RAGE. An indication of oxidative damage has been proposed to be the irreversible oxidative alteration of proteins known as protein carbonylation. Our data support Dhivya et al.’s (2017) findings that proteins were carbonylated as a result of the oxidative stress generated by isoproterenol [55]. As a result, it implies that rats’ ISO-induced cardiac damage may be caused by oxidative stress spurred by advanced glycation end products. The use of rosuvastatin, however, prevented cellular damage by raising antioxidant activities, lowering lipid peroxidation, and protecting proteins, which together lowered ISO-induced oxidative stress. The initial line of cellular defence against oxidative stress has been suggested to consist of SOD, CAT, GST, GPx, and GR. In our investigation, cardiac activity levels of rats receiving ISO treatment were shown to be considerably lower than those of control rats for SOD, CAT, GST, GPx, and GR. The activities may have decreased as a result of their inactivation and increasing use in ROS scavenging. Myocardial cell membranes are more sensitive to oxidative damage as a result of their diminished activity. The cell’s endogenous antioxidant defence system, which consists of antioxidant enzymes and non-enzymatic substances, keeps the redox equilibrium in balance. As it takes into account the cumulative antioxidant effect within a sample, this shows the overall antioxidant activity of a system. The Trolox equivalents in tissue homogenates from the ISO group were noticeably lower. These outcomes are consistent with Kocak et al.’s (2016)[56] findings. In comparison to the ISO group, pretreatment with rosuvastatin stabilised TAA at a greater level. These outcomes support the conclusions drawn from the study’s oxidant and antioxidant parameters.

In the current investigation, AGEs induction considerably raised serum AGEs levels and AGEs accumulation. Circulating AGE concentrations are correlated with negative clinical outcomes and the severity of coronary artery disease [57]. Although the precise mechanisms are unknown, rosuvastatin can lower the serum level of AGEs in a way that is both time-dependent and independent of decreasing cholesterol. A source of free-radical superoxide generation on their own, AGEs are formed as a result of oxidative stress [58, 59]. Additionally, it has been demonstrated that the hydroxyl metabolites of rosuvastatin have anti-oxidative characteristics [60].

We looked into whether statins prevent VSMC proliferation by preventing ROS-driven proliferation and the inflammatory pathway as ROS is known to be generated via the activation of NADPH oxidase from the interaction between AGE and RAGE. This theory is based on the concept that the AGE-RAGE interaction may be the upstream mechanism that enhances ROS production and the subsequent proliferation of VSMC.

Rosuvastatin has been shown to have antioxidant properties in recent experimental experiments, and it may also reduce ROS in endothelial cells by S-nitrosylating thioredoxin [61,62]. Recent investigations have raised the possibility that rosuvastatin may function as an antioxidant in rabbit models of experimental atherosclerosis [63,64], preventing the oxidation of LDL by activated macrophages generated from monocytes. As in other investigations, the ROS production in our data rose with the activation of AGEs and decreased with statin administration.

In the current study, rosuvastatin decreased enhanced intracellular oxidative stress (ROS) spurred by the AGE-RAGE interaction as well as ROS-induced cellular signalling.

According to the results of the current investigation, rosuvastatin dramatically reduced the expression of HMGB1 and RAGE. Rosuvastatin inhibited RAGE expression in rat aortas, according to studies by Xu et al [65], Yang et al [11], Jin et al [66], and Yang et al. Rosuvastatin also reduced HMGB1 activation and serum levels, according to studies by Yang et al. and Jin et al. TNF-α and IL-6 are two examples of the proinflammatory cytokines that are upregulated as a result of HMGB1 binding to RAGE [67]. These findings are corroborated by a number of earlier investigations [69] as well as Gao et al. (68), who demonstrated that HMGB1 and RAGE mediated the overexpression of TNF-α.

Additionally, after an inflammatory damage, HMGB1 binding to RAGE causes an upregulation of anti-inflammatory cytokines such IL 4 and IL 10 [69–71]. Rosuvastatin post-conditioning lowered oxidative stress indicators in rats after ischemia/reperfusion injury, according to research by Du et al. (48). Furthermore, Ke et al. [47] demonstrated that rosuvastatin preconditioning reduced ischemia/reperfusion injury by reducing the accumulation of inflammatory cells in the heart, which is connected to increased production of inflammatory cytokines, such as TNF α and IFN-γ, and dysregulated anti-inflammatory cytokines, such as IL 4 and IL-10. The delivery of rosuvastatin to the ISG group led to a rise in TNF-α and IL-6 levels as well as the expression of HMGB1 and RAGE, according to the results of the current investigation. All of these findings imply that rosuvastatin’s stimulation of HMGB1/RAGE may lead to a better functional recovery. However, more research is required to determine how rosuvastatin affects inflammatory cells.

The current study’s findings showed that rosuvastatin significantly improved left ventricular structure and function, also known as LVEF. This may be due to a number of factors, including how long rosuvastatin is given for. According to findings that claim rosuvastatin’s potential to improve LVEF is time-dependent [72], long-term therapy may be more beneficial in enhancing LVEF. Before affecting cardiac tissue, rosuvastatin was thought to affect inflammatory cytokine expression and serological markers. More research is needed to solve this issue.

Mitochondria are important subcellular organelles that play a role in energy metabolism and are susceptible to oxidative damage. Mitochondria, which are prevalent in cardiomyocytes, are the main organelles linked to cardiac damage [73]. This may have an impact on both the origin and outcome of oxidative heart injury. According to the literature, AGE-RAGE causes the mitochondrial respiratory chain to behave differently by downregulating particular mitochondrial respiratory enzyme functions, decreasing antioxidant potential, increasing the production of mitochondrial ROS, and promoting necrosis in the heart [74].

The effects of TCA cycle enzymes or mitochondrial respiratory chain enzymes including IDH, KDH, SDH, and MDH were considerably reduced in the ISO-treated rats when compared to NCG rats in the current study. ROS may impede the actions of these enzymes, which could have an effect on the oxidation of the substrate in the mitochondria, the reduction of action transfer rates that are equal to molecular oxygen molecules, and the catabolism of cellular energy [75]. When compared to the ISO group, rosuvastatin treatment significantly raised the number of enzymes. When compared to an NCG, the per se group demonstrated an increase in the number of mitochondrial enzymes in the myocardium.

Collagen is the protein that is most common in the extracellular cardiovascular matrix. Collagen fibres in the heart create a network of linking myocytes that help to preserve the shape of the ventricles and transmit the contractile force from all myocardial cells to the ventricular canal. preserving the cardiac tissue’s structure is essential for preserving the shape and functionality of the heart chambers. The heart contains both I and III forms of collagen. Collagen I in particular has a considerable effect on ventricular stiffness due to its tough and tensile properties [76,77].

## 5. Conclusion

By showing that the functional AGE/RAGE axis is inhibited following rosuvastatin medication, this study concludes by putting out an etiologic hypothesis regarding the role of rosuvastatin therapy in heart injury. These findings may also have practical significance since they highlight the intriguing possibility that rosuvastatin’s alteration of AGE-RAGE signalling may offer a cutting-edge treatment option for reducing the onset and progression of other cardiovascular diseases.

This study demonstrates the clinical relationship between the AGE-RAGE axis and heart injury. This study also highlights rosuvastatin’s preventive effects against heart diseases. According to the study, rosuvastatin has cardioprotective effects on the experimental model, which were supported by a number of physical, biochemical, and histological characteristics. Future research should be conducted to identify the precise molecular mechanisms and signalling pathways of atorvastatin’s impact on the production of AGEs-RAGE.

## Author Contributions

Conceptualization, R.W., T.M., M.H.S; methodology, R.W., T.M. and A.S.; software, T.M., R.W. and F.A.; validation, T.M. and M.H.S; formal analysis, T.A.W. and S.Z.; investigation, T.M. and A.S.; resources, T.M. and R.W.; data curation, R.W., M.S. and R.W.; writing—original draft preparation, M.S. and S.P.; writing—review and editing, A.S. and T.A.W.; visualization, R.W..; supervision, T.M; M.W. and M.H.S; project administration, T.M.; funding acquisition, R.W. and T.A.W. All authors have read and agreed to the published version of the manuscript.

## Institutional Review Board Statement

The Institutional Animal Ethics Committee (IAEC) of the Faculty of Pharmacy, Integral University, Lucknow (U.P.), India, approved the study process with approval number (IU/IAEC/21/09), (Reg no. 1213/PO/Re/S/08/CPCSEA, 5 June 2008).

## Informed Consent Statement

Not applicable.

## Data Availability Statement

Data are contained within the article.

## Acknowledgments

The authors are grateful to Integral University’s Honorable Founder and Chancellor, Syed Waseem Akhtar, and Vice-Chancellor, Javed Musarrat, for providing an exceptional research atmosphere and resources. The authors extend their appreciation to the Researchers Supporting Project Number (RSP2023R357). King Saud University, Riyadh, Saudi Arabia, for funding this work. (IU/R&D/2022-MCN0001655).

